# Spatio-temporal diversity of dietary preferences and stress sensibilities of early and middle Miocene Rhinocerotidae from Eurasia: Impact of climate changes

**DOI:** 10.1101/2022.05.06.490903

**Authors:** Manon Hullot, Gildas Merceron, Pierre-Olivier Antoine

## Abstract

Major climatic and ecological changes are documented in terrestrial ecosystems during the Miocene epoch. The Rhinocerotidae are a very interesting clade to investigate the impact of these changes on ecology, as they are abundant and diverse in the fossil record throughout the Miocene. Here, we explored the spatio-temporal evolution of rhinocerotids’ paleoecology during the early and middle Miocene of Europe and Pakistan. We studied the dental texture microwear (proxy for diet) and enamel hypoplasia (stress indicator) of 19 species belonging to four sub-tribes and an unnamed clade of Rhinocerotidae, and coming from nine Eurasian localities ranging from Mammal Neogene zone (MN) 2 to MN7/8. Our results suggest clear differences in the feeding ecology and thus niche partitioning at Kumbi 4 (MN2, Pakistan), Sansan (MN6, France), and Villefranche d’Astarac (MN7/8, France), while overlap of the interpreted diets and subtle variations are discussed for Béon 1 (MN4, France) and Gračanica (MN5/6, Bosnia-Herzegovina). All rhinocerotids studied were interpreted as browsers or mixed-feeders, and none had a grazer nor frugivore diet. The prevalence of hypoplasia was moderate (*∼*10%) to high (> 20%) at all localities but Kumbi 4 (*∼*6%), and documented quite well the local conditions. For instance, the high prevalence at the close to Miocene Climatic Optimum locality of Béon 1 (*∼*26%) has been correlated with periodical droughts, while the moderate ones (*∼*10%) at Sansan and Devínska Nová Ves Spalte (Slovakia) both dated from the MN6 (i.e., by the middle Miocene Climatic Transition, ca. 13.9 Mya) were linked to the persistence of sub-tropical local conditions. Besides locality, species and tooth locus were also important factors of variation for the prevalence of hypoplasia. The very large hippo-like *Brachypotherium brachypus* was one of the most affected species at all concerned localities (but Sansan), whereas early-diverging elasmotheriines were very little affected, suggesting an influence of phylogeny and/or diet in stress susceptibility.

## Introduction

The Miocene is a key period in Earth and rhinocerotid evolutionary histories. Climatic conditions in Eurasia during the Miocene epoch are globally warm and the typical habitat is forested (Cerling et al., 1997; Zachos et al., 2001; Bruch et al., 2007; Westerhold et al., 2020). It is the last warm episode of the Cenozoic era, although marked by great climatic changes prefiguring the setup of modern cold conditions (Westerhold et al., 2020). During early Miocene times, temperatures increased until reaching the Miocene Climatic Optimum (MCO) between ∼17 to 14 Mya (corresponding to the late Burdigalian + Langhian standard ages; Westerhold et al., 2020). This optimum is followed by an abrupt cooling (the middle Miocene climatic transition [mMCT]; Westerhold et al., 2020) associated with faunal turnovers in Eurasia (Böhme, 2003; Maridet et al., 2007). The middle Miocene is marked by a relative aridity, associated with a global cooling (Bruch et al., 2007; Böhme et al., 2008).

Regarding rhinocerotids, Miocene times witness peaks in their alpha diversity about 22–18 Mya and 11–10 Mya (Antoine et al., 2010; Antoine and Becker, 2013; Antoine, in press.). During the early and middle Miocene in Eurasia, four sub-tribes of Rhinocerotidae are encountered – Rhinocerotina, Teleoceratina and Aceratheriina (Rhinocerotinae), and Elasmotheriina (Elasmotheriinae) – and species of which are often found associated in fossil-yielding localities (Antoine et al., 1997, 2010; Heissig, 2012; Becker and Tissier, 2020; Antoine, 2002, in press.). This abundance and the potential cohabitation of such large herbivores question habitat capacity and competition for food resources. However, the ecology of the rhinocerotids has rarely been explored (e.g., Mihlbachler et al., 2018; Wang and Secord, 2019; Rivals et al., 2020; Stefaniak et al., 2020) or only been assumed based on morphological adaptations (as criticized by Prothero et al., 1989; Prothero, 2005; Giaourtsakis et al., 2006). If the Rhinocerotina appear to be ecologically varied, the literature suggests a similar ecology for most representatives of certain tribes. For instance, the elasmotheriines are considered as open environment dwellers adapted to though vegetation (Iñigo and Cerdeño, 1997; Antoine and Welcomme, 2000), whereas the teleoceratines are reconstructed as hippo-like rhinoceroses inhabiting lake side or swamps and probably browsing on low vegetation or even grazing (Prothero et al., 1989; Cerdeño, 1998), although a semi-aquatic lifestyle similar to hippos is debated (e.g., Clementz, 2012). With this article, we wanted to assess if the great diversity of Miocene Rhinocerotidae was indeed associated to an ecological diversity and to explore paleoecological differences associated with climate changes between the studied species, regions, and periods. To do so, we focused on the rhinocerotids from nine localities, covering wide temporal and geographical ranges (from early to middle Miocene, Mammal Neogene zone [MN] 2 to MN7/8, and from southwestern France to Pakistan). We coupled dental microwear texture analysis (short-term diet proxy) and enamel hypoplasia (i.e., enamel defect resulting from a stress stopping tooth development) to infer dietary preferences and stress sensibility, respectively. Eventually, we compared the results of both approaches, which has the potential to detect stressful shifts in diet (competition, punctual shortage event) or accentuated susceptibility due to a specialized diet, and put them into phylogenetic and environmental contexts.

## Material and methods

We studied the rhinocerotid dental remains from nine early and middle Miocene localities from France (Béon 1, Béon 2, Sansan, Simorre, and Villefranche d’Astarac), Germany (Steinheim am Albuch), Bosnia-Herzegovina (Gračanica), Slovakia (Devínska Nová Ves Spalte), and Pakistan (Kumbi 4, Balochistan), ranging from MN2 to MN7/8. The rhinocerotid assemblages are detailed in Table 1. The specimens are curated at the Naturhistorisches Museum Wien (NHMW), the Muséum de Toulouse (MHNT), and the Naturhistorisches Museum Basel (NHMB). For all details on the specimens included in this study see supplementary S1. The localization of the studied localities is given in Figure 1. Further details on the localities are given in supplementary S2.

**Figure 1:**
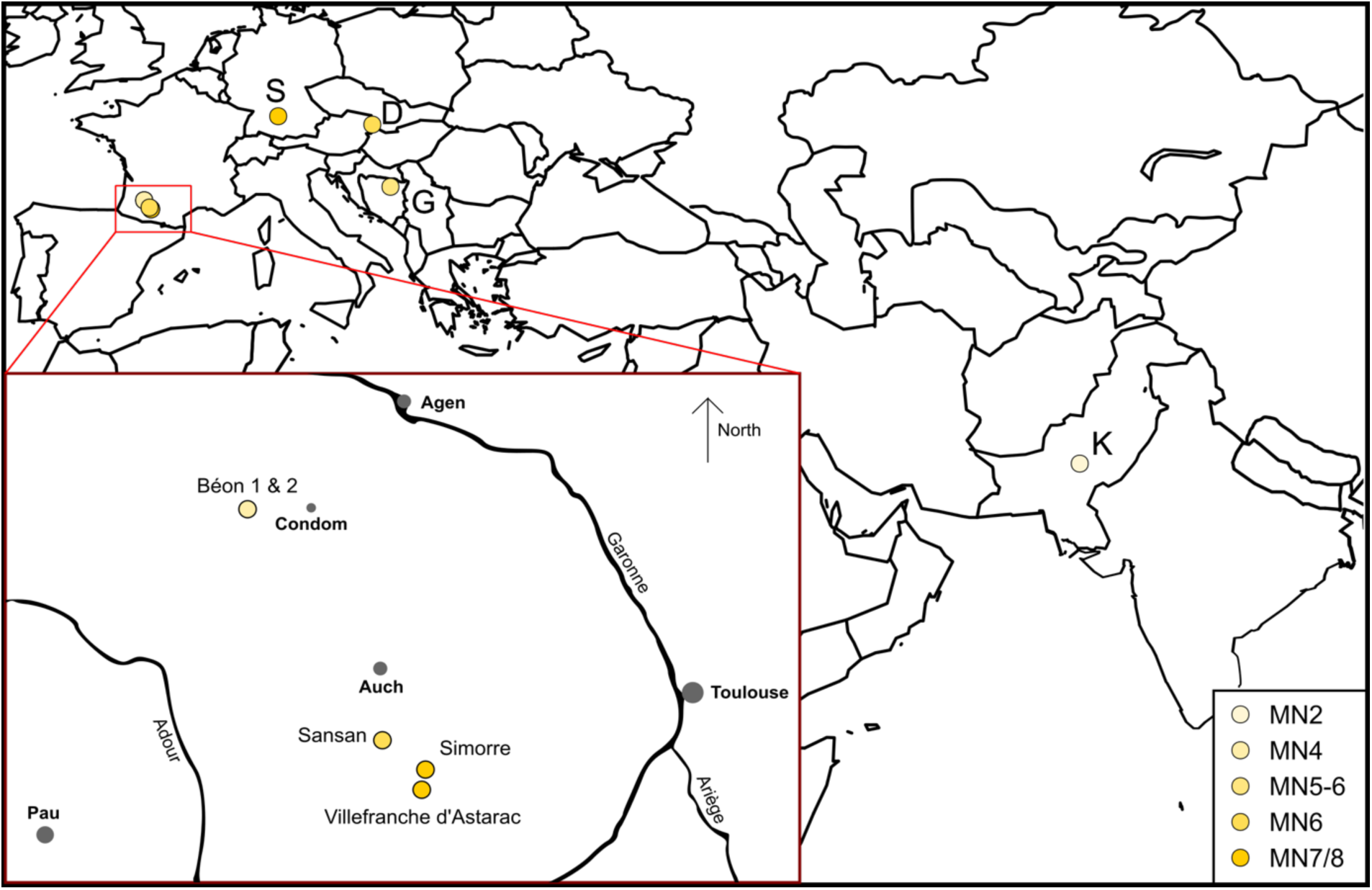
Geographical position of the studied Eurasian Miocene localities. Localization of all localities in Eurasia. Red square is a zoom on the southwestern French localities, modified from Antoine and Duranthon (1997). Color code by MN zones as detailed on the lower right corner. Abbreviations from West to East: S-Steinheim am Albuch (MN7/8; Germany), D-Devínska Nová Ves Spalte (MN6; Slovakia), G-Gračanica (MN5-6; Bosnia-Herzegovina), K-Kumbi 4 (MN2; Pakistan).

**Table 1:**
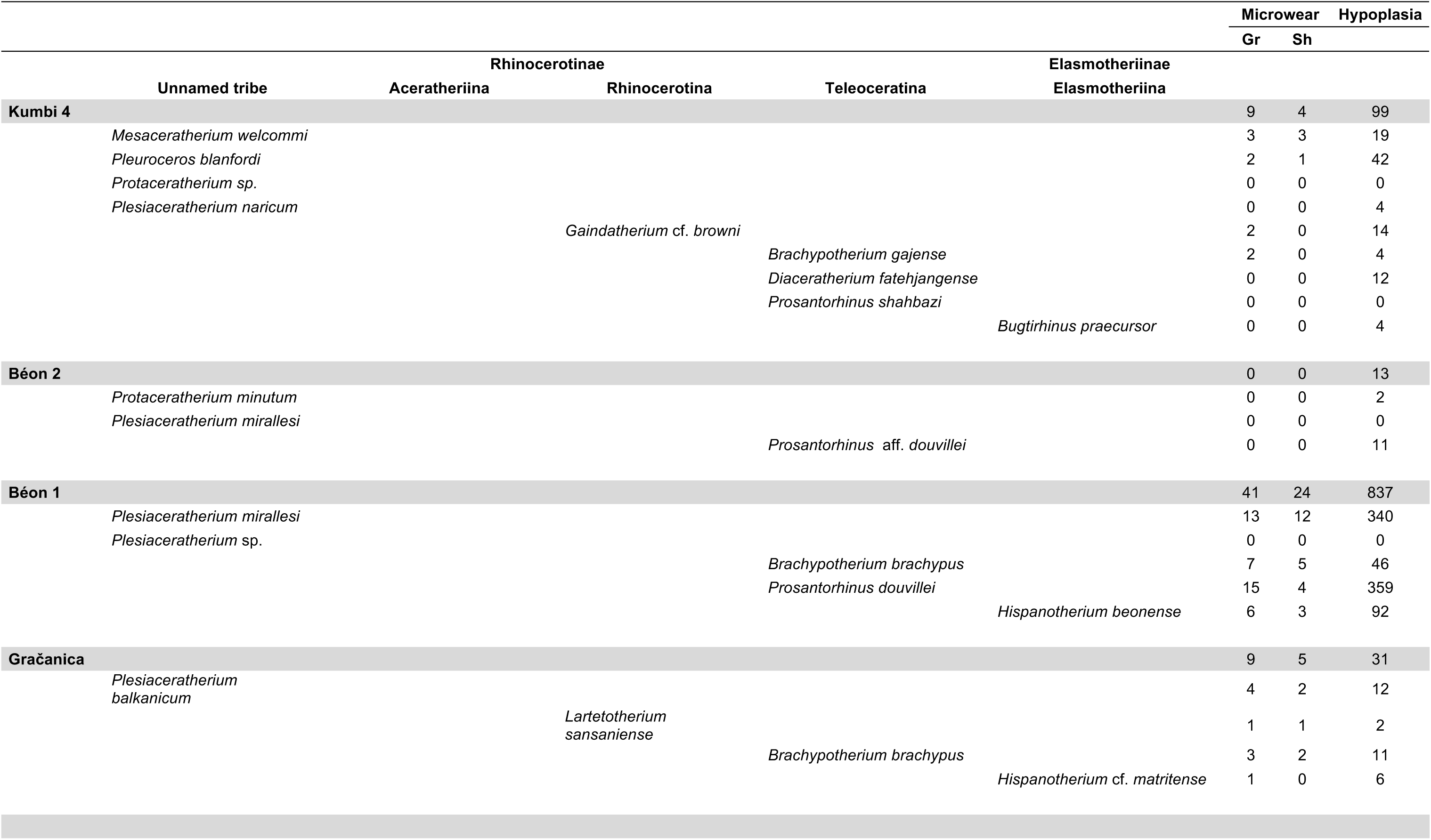
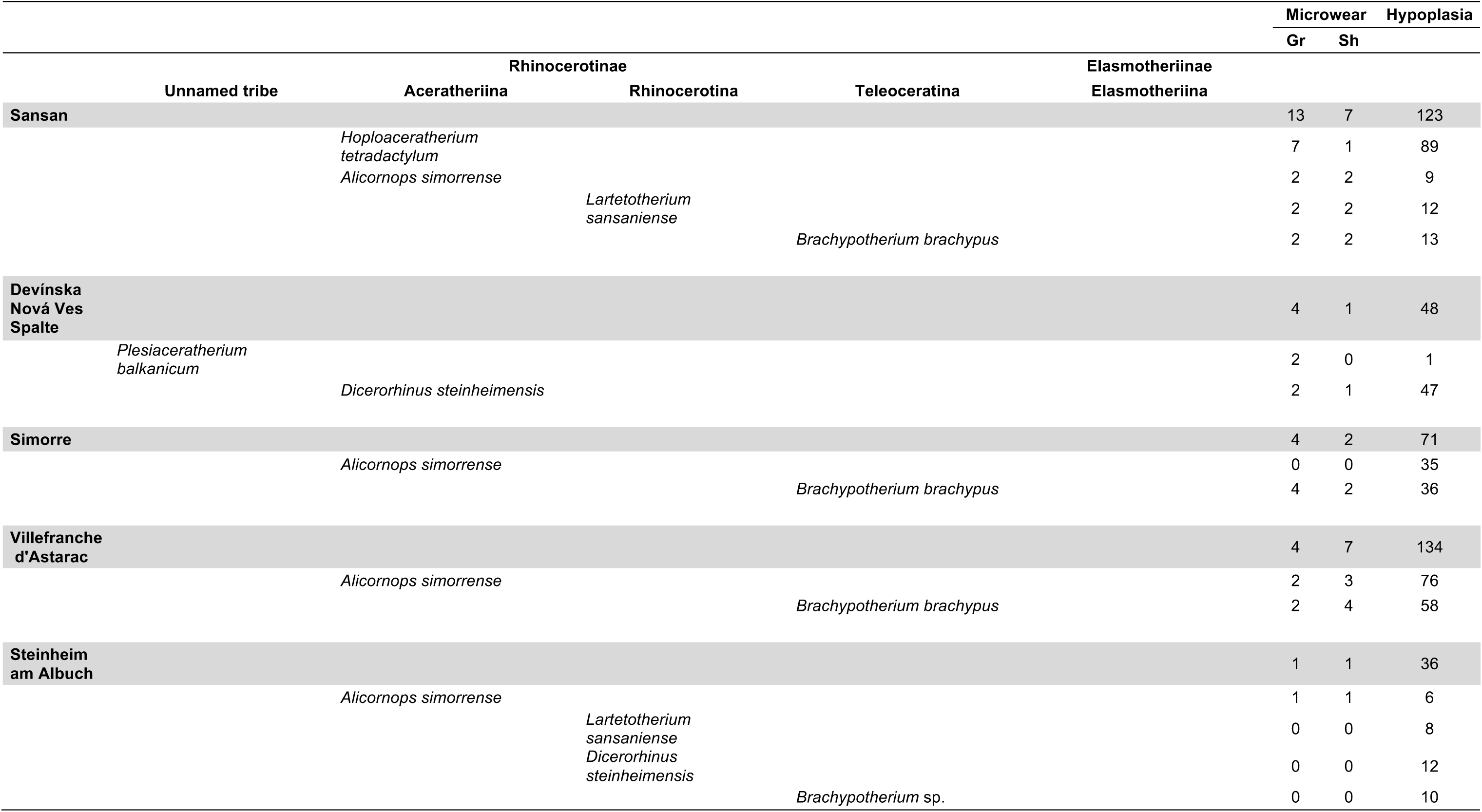
List of rhinocerotid species found at each locality studied along with the number of specimens included for each method.

### Dental Microwear Texture Analyses (DMTA)

Dental Microwear Texture Analysis (DMTA) allows for the characterization of dietary preferences at a short time scale (days to weeks prior the death of the individual; Hoffman et al., 2015; Winkler et al., 2020). This method has been widely used in paleontological and archeological studies to infer diet (e.g., Grine, 1986; Rivals et al., 2012; Jones and DeSantis, 2017; Berlioz et al., 2018). Here, we studied dental microwear texture on one well-preserved molar (germs and over-worn teeth excluded) per individual, preferentially the second molar (first or third otherwise), either upper or lower, left or right. Indeed, the choice of the locus and the facet are crucial in microwear analyses, as differences have already been noted and may impact the interpretations (Mihlbachler et al., 2016; Ramdarshan et al., 2017).

After cleaning the tooth with acetone or ethanol, two silicone (Regular Body President, ref. 6015 - ISO 4823, medium consistency, polyvinylsiloxane addition type; Coltene Whaledent) molds were made on a single enamel band, which shows two different facets acting as grinding and shearing (if present). The grinding facet shows several Hunter-Schreger bands on the very enamel surface – due to the fact that they are vertical in Rhinocerotoidea (Koenigswald et al., 2011) – and is more horizontal than the shearing facet which has a steep slope towards the labial side (Butler, 1952). To combine both types of facets with different functions indeed improves dietary reconstruction (Louail et al., 2021; Merceron et al., 2021). The enamel band on which we identified those two facets is localized labially near the protocone on upper molars and distally to the protoconid or hypoconid (if the protoconid is unavailable) on lower teeth (for an illustration see supplementary S2).

In this article we followed a protocol adapted from Scott et al. (2005, 2006) with sensitive-scale fractal analyses. Molds were scanned with a Leica DCM8 confocal profilometer (“TRIDENT” profilometer housed at the PALEVOPRIM, CNRS, University of Poitiers) using white light confocal technology with a 100× objective (Leica Microsystems; Numerical aperture: 0.90; working distance: 0.9 mm). The obtained scans (.plu files) were pre-treated with LeicaMap v.8.2. (Leica Microsystems) as follows: the surface was inverted (as scans were made on negative replicas), missing points (i.e., non-measured, less than 1%) were replaced by the mean of the neighboring points and aberrant peaks were removed (see details in the supplementary information in Merceron et al., 2016). The surface was then levelled, and we applied a polynomial of degree 8 removal of form to temper for Hunter-Schreger bands reliefs in the DMTA parameters. Eventually, we selected a 200×200-μm area (1551 × 1551 pixels) within the surface, which we saved as a digital elevation model (.sur) and used to extract DMTA parameters through Scale-Sensitive Fractal Analysis with SFrax (Surfract, www.surfract.com) and LeicaMap.

Here we focused on five classical DMTA parameters: anisotropy (exact proportion of length-scale anisotropy of relief; epLsar), complexity (area-scale fractal complexity; Asfc), heterogeneity of complexity (heterogeneity of area-scale fractal complexity here at 3×3 and 9×9; HAsfc9 and HAsfc81), and fine textural fill volume (here at 0.2 μm; FTfv). The description of these parameters is available in Scott et al. (2006).

To facilitate DMTA interpretation for fossil specimens, we used the values and thresholds proposed in extant species by Scott (2012) and Hullot et al. (2019). For the latter dataset, two new specimens were added to increase sample size (notably for the Indian rhinoceros, which has a diet changing with the season), as specified thereafter, and consists of 17 specimens of *Ceratotherium simum* (white rhinoceros), four of *Dicerorhinus sumatrensis* (Sumatran rhinoceros), 21 of *Diceros bicornis* (black rhinoceros), 15 of *Rhinoceros sondaicus* (Javan rhinoceros; one new specimen), and five of *Rhinoceros unicornis* (Indian rhinoceros; one new specimen).

### Enamel hypoplasia

Hypoplasia is a common defect of the enamel resulting from a stress or a combination of stresses occurring during tooth development (Goodman and Rose, 1990). It is a permanent, sensitive, but non-specific indicator of stresses either environmental (e.g., drought or nutritional stress; Skinner and Pruetz, 2012; Upex and Dobney, 2012), physiological (e.g., disease or parasitism; Suckling et al., 1986; Rothschild et al., 2001; Niven et al., 2004), and/or psychological (e.g., depression in primates; Guatelli-Steinberg, 2001).

Enamel hypoplasia was studied with the naked eye and categorization of the defects followed the *Fédération Dentaire Internationale* (1982) as linear enamel hypoplasia (LEH), pitted hypoplasia, or aplasia. We studied all cheek teeth, both deciduous and permanent, but excluded 62 teeth to avoid false negative and uncalibrated defects, as enamel was obscured (e.g., tooth unerupted in bone, sediment occluding), broken or worn out, or as identification of the locus was impossible. This left 1401 teeth studied for the hypoplasia analysis – 294 milk molars and 1107 permanent premolars and molars – from the nine localities. In parallel, qualitative data (tooth locus affected, position of the defect on the crown, and severity) and caliper measurements (distance of the defect from enamel-dentine junction, width if applicable) were taken (details in supplementary S3). This data may give insights into the age at which the defect occurred (locus affected, position on the crown, distance to enamel-dentine junction), as well as into the duration and intensity of the stress (width of the defect), which can in turn help us propose causes (e.g., birth for a defect near the base of the crown on a D4). Type of defects recorded, and caliper measurements are illustrated in Figure 2.

**Figure 2:**
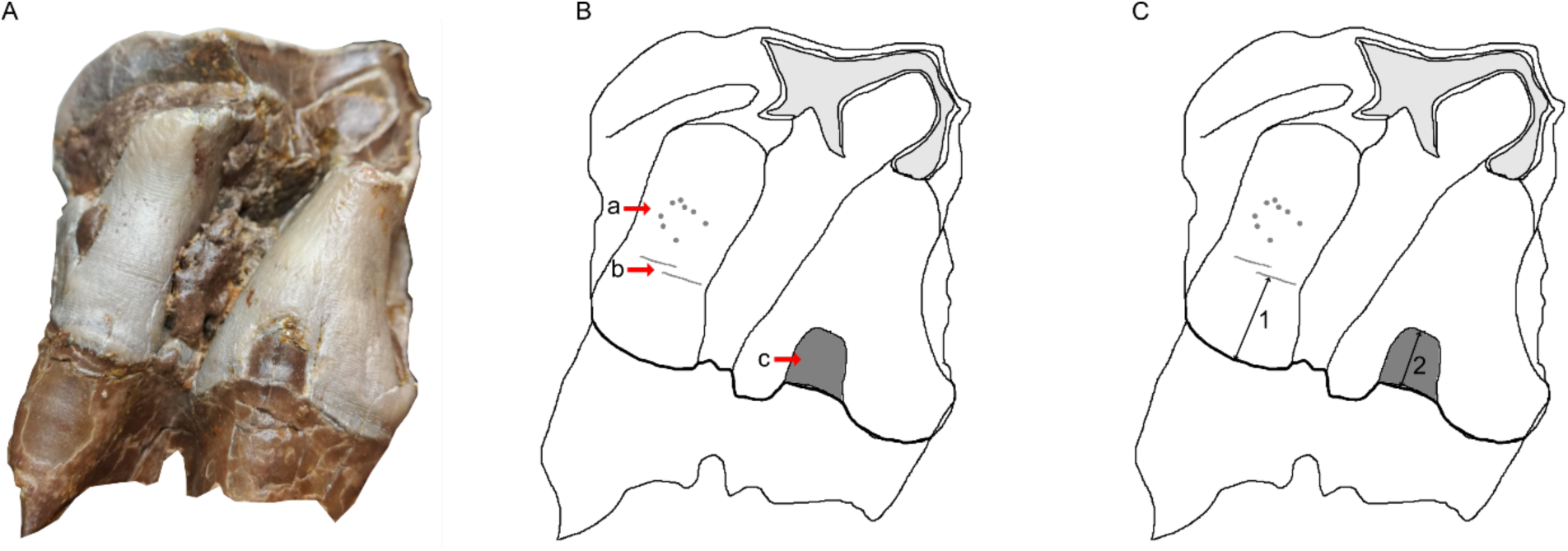
The three different types of hypoplasia considered in this study and the associated measurements. A-Lingual view of right M2 of the specimen MHNT.PAL.2004.0.58 (*Hispanotherium beonense*; Béon 1) displaying three types of hypoplasia B-Interpretative drawing of the photo in A illustrating the hypoplastic defects: a-pitted hypoplasia, b-linear enamel hypoplasia, and c-aplasia C-Interpretative drawing of the photo in A illustrating the measurements: 1-distance between the base of the defect and the enamel-dentin junction, 2-width of the defect (when applicable). Figure from Hullot et al. (2021).

### Statistics and GLMMs

Statistics were conducted in R (R Core Team, 2018: https://www.R-project.org/), equipped with the following packages: reshape2 (Wickham, 2007), dplyr (Wickham et al., 2019), lme4 (Bates et al., 2015), car (Fox et al., 2012), MASS (Venables and Ripley, 2002). According to the recent statement of the American Statistical Association (ASA) on p-values (Wasserstein and Lazar, 2016; Wasserstein et al., 2019), we avoided as much as possible the use of the term “statistically significant” associated to the classical thresholds in this manuscript. However, some typical thresholds have still been used for sake of clarity. Figures were done using R package ggplot2 (Wickham, 2011) as well as Inkscape v.0.91.

General Linear Mixed Models (GLMM) on our data were constructed based on a R code modified from Arman et al. (2019) and adapted to each tested response variable. An example of this code applied to hypoplasia variable Hypo is given in supplementary S4. DMTA response variables were the five DMTA parameters (epLsar, Asfc, FTfv, HAsfc9, and Hasfc81) and we selected Gaussian family for the GLMMs. Factors in the models were: specimen (number of the specimen; random factor), locality, province, age (MN zones), genus, tooth (e.g., second molar, fourth milk molar), position (upper or lower), side (left or right), cusp (protocone, protoconid, hypoconid), and facet (grinding or shearing). For hypoplasia, response variables were Hypo (1 or 0 for presence or absence of hypoplasia, respectively) for which we used Binomial family, Defect (e.g., LEH, Pits, Aplasia; converted to numbers), Localization (position of the defect on the crown; mostly labial or lingual, converted to numbers), Multiple (number of defects), and Severity (0 to 4), modeled using Poisson family. The factors were: specimen (number of the specimen; random factor), locality, province, age (MN zones), genus, tooth (e.g., first molar, fourth premolar), position (upper or lower), side (left or right), and wear (low, average, high). Additionally, for response variables Severity, Multiple, and Localization, defect was converted and used as a factor.

The models were built with a bottom-up approach, starting with the only random factor of our dataset alone (specimen) and adding factors incrementally for every set (e.g., 1|Specimen + Genus, 1|Specimen + Locality). New set was built as long as Akaike’s Information Criterion score (AIC) kept decreasing. Few interactions (e.g., Genus x Facet for microwear, Genus x Tooth for hypoplasia) were tested in the models, as most factors were considered independent and to avoid unnecessarily complex and rarely selected models (Arman et al., 2019). We selected the best candidate model as the one with the lowest AIC and checked for over-dispersion (ratio of deviance and degrees of freedom > 1.5). If needed (over-dispersion), we corrected it through quasi-Poisson or quasi-Binomial laws from the MASS package (Venables and Ripley, 2002) or by adjusting the coefficients table (multiply type error by square root of the dispersion factor and recalculate Z and p values accordingly). In total, 340 models were compared across the 10 response variables (see supplementary S5, S6, and S7).

## Results

### Microwear

MANOVA (Species x Facet x Age x Locality) on all five main DMTA parameters (epLsar, Asfc, FTfv, Hasfc9, Hasfc81) revealed low p-values for Species (df = 14; p-value = 8.6 x 10^-4^), Facet (df = 1; p-value = 6.5 × 10^-4^), and Locality (df = 4; p-value = 0.014). However, when we separated our sample by facet, most of these differences were only found in the shearing facet subsample (n = 51), while very few were observed for the grinding facet subsample (n = 85). Based on MANOVA results, ANOVAs (Species x Age x Locality) were conducted for each parameter in each microwear facet subsample. In the grinding subsample, we only found a potential influence of Species (p-value = 0.0165) and Age (p-value = 0.0426) on Asfc. In the shearing subsamples, Species had an impact on epLsar (p-value = 0.034) and Hasfc 9 (p-value = 0.0477), while Age had an impact on Asfc (p-value = 4.31 × 10^-4^) and FTfv (p-value = 0.035).

As the different factors (Species and Age) had more than two states, we conducted post hoc tests to precise the highlighted differences, results of which are detailed in Table 2 and Table 3. The more conservative post hoc (Tukey’s honestly significant difference; HSD) revealed very few noticeable differences in the microwear textures of the studied rhinocerotid specimens (Table 2). On the shearing facet *Plesiaceratherium mirallesi* (n = 12) had higher values of epLsar than *Brachypotherium brachypus* (n = 15; p-value = 8.81 × 10^-3^), *Mesaceratherium welcommi* (n = 3; p-value = 0.040), and *Lartetotherium sansaniense* (n = 3; p-value = 0.016). The shearing facet of the rhinocerotids from the MN5 (i.e., Gračanica, n = 5) had lower values of Asfc than that from the MN6 (Sansan and Devínska Nová Ves, n = 8; p-value = 0.058) and MN7/8 (Simorre, Villefranche and Steinheim, n = 10; p-value = 0.056). The least conservative post hoc (Fischer’s least significant difference; LSD) highlighted more differences in the DMTA patterns of the specimens regarding Species and Age (Table 3, p-values < 0.05). Concerning Species, *P. mirallesi* (n = 12) and *A simorrense* (n = 5) had higher values of epLsar on their shearing facet compared to *B. brachypus* (n = 15), *P. balkanicum* (n = 2), *P. douvillei* (n = 4), *tetradactylum* (n = 1), *L. sansaniense* (n = 3), *D. steinheimensis* (n = 1), and *M. welcommi* (n = 3). Regarding Asfc, *D. gajense* (n = 2), *G.* cf. *browni* (n = 2) and *H. tetradactylum* (n = 7) had higher values compared to *P. douvillei* (n = 15), *A. simorrense* (n = 4), *H.beonense* (n = 6), *P. mirallesi* (n = 13), *H.* cf. *matritense* (n = 1). However, with the p-value threshold of 0.05 selected for LSD tests, no differences in Hasfc9 were found by Species. Concerning Age, MN4 (Béon 1) and MN5 (Gračanica) rhinocerotids had lower complexity values than those from MN6 (Sansan and Devínska Nová Ves) and MN7/8 (Simorre, Villefranche and Steinheim) for both facets, and MN2 (Kumbi 4) for the grinding facet only. The FTfv was also different on the shearing facet of rhinocerotids from MN2 (n = 4) and from MN4 (n = 24), MN5 (n = 5), and MN6 (n = 8), and between the rhinocerotids from MN5 and those from MN7/8 (n = 10).

**Table 2:** Pairs (Species or Age) with low p-values after Tukey’s honestly significant difference (HSD) by DMTA parameters. FTfv not shown as it yielded p-values above 0.1 only

| DMTA parameter | Facet | Pair (Species or Age) with differences |  | p-value |
| --- | --- | --- | --- | --- |
| **Asfc** | **Grinding** | *Hoploaceratherium tetradactylum* | *Plesiaceratherium mirallesi* | 0.034 |
|  |  |  | *Prosantorhinus douvillei* | 0.055 |
|  | **Shearing** | MN5 | MN6 | 0.058 |
|  |  |  | MN7/8 | 0.056 |
| **epLsar** | **Shearing** | *Plesiaceratherium mirallesi* | *Brachypotherium brachypus* | 0.009 |
|  |  |  | *Lartetotherium sansaniense* | 0.016 |
|  |  |  | *Mesaceratherium welcommi* | 0.040 |

**Table 3:**
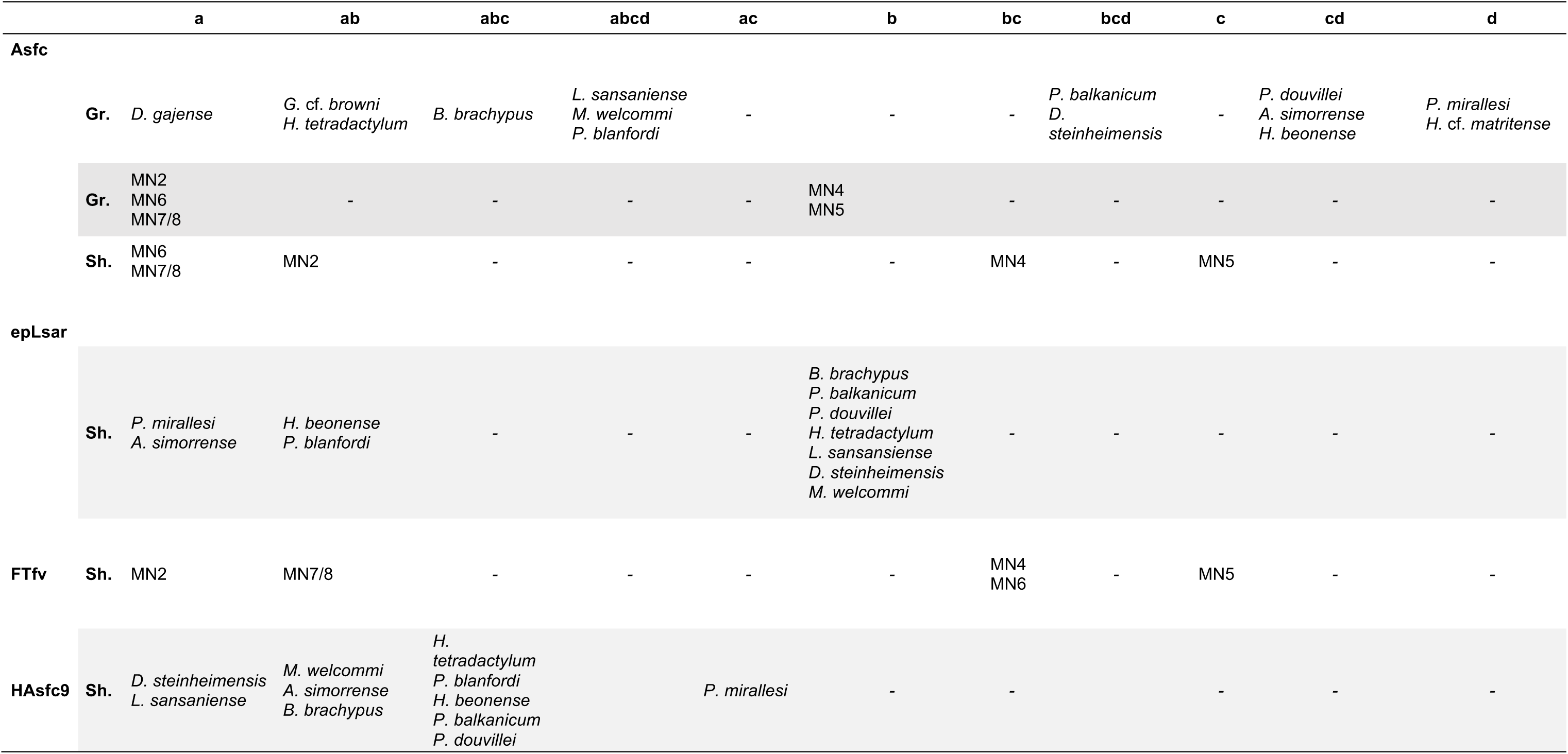
Fischer’s least significant difference (LSD) post hoc results by DMTA parameters. Groups (a, ab, abc, abcd, ac, b, bc, bcd, c, cd, d) are significantly different from one another (here with a 0.05 threshold, as a p-value threshold had to be defined to run the LSD tests) if they don’t share a common letter (e.g., ab is distinct from c, cd, and d). Gr. and Sh. stands respectively for grinding and shearing facets. No post hocs conducted for HAsfc81 as ANOVA did not reveal differences for this parameter.

Except at Béon 1, the specific microwear sampling at each locality was very restricted (n < 5), either due to low numbers of exploitable teeth available, or to the lack of well-preserved microwear texture on molars. In order to facilitate the understanding, the results are presented by locality (chronologically) and by species. All dietary interpretations in the results section are based on reference dataset and values in extant species of bovids (Scott 2012) and rhinoceros (Hullot et al. 2019).

At **Kumbi 4**, four species were considered for DMTA: *Pleuroceros blanfordi*, *Mesaceratherium welcommi*, *Gaindatherium* cf. *browni* (grinding facet only), and *Brachypotherium gajense* (grinding facet only). Figure 3 shows that Kumbi rhinocerotids display a great variety of microwear patterns. Only one specimen, belonging to *G.* cf. *browni*, is above the high anisotropy threshold of 5 x 10^-3^, while regarding complexity all specimens of *B. gajense* and *G.* cf. *browni* but none of *P. blanfordi* display values above the high complexity cutpoint of 2. *Gaindatherium* cf. *browni* and *P. blanfordi* have large variations in anisotropy, from low values (∼ 1 x 10^-3^) to high (about 5 x 10^-3^), but consistent values of complexity (around 3 and 1.4 respectively). Such a pattern associated with moderate (HAsfc9: 0.3-0.4, HAsfc81: 0.5-0.6; *P. blanfordi*) to high (HAsfc9 > 0.3, HAsfc81 > 0.6; *G.* cf. *browni*) values of HAsfc (Figure 4) point towards mixed-feeding diets, probably with the inclusion of harder objects for *G.* cf. *browni* (higher values of complexity). The signature for *B. gajense* is suggestive of browsing with low mean anisotropy (1.82 x 10^-3^), but high means of complexity (4.58), FTfv (7.89 x10^4^), and HAsfc (HAsfc9 = 0.36; HAsfc81 = 1). Eventually, *M. welcommi* presents low to moderate anisotropy values (< 4 x 10^-3^), moderate complexity (∼ 1.5) and HAsfc (HAsfc9: 0.2-0.4, HAsfc81: 0.3-0.6), but high FTfv (> 4 x 10^4^) on both facets (Figure 4), which denotes browsing or mixed-feeding habits.

**Figure 3:**
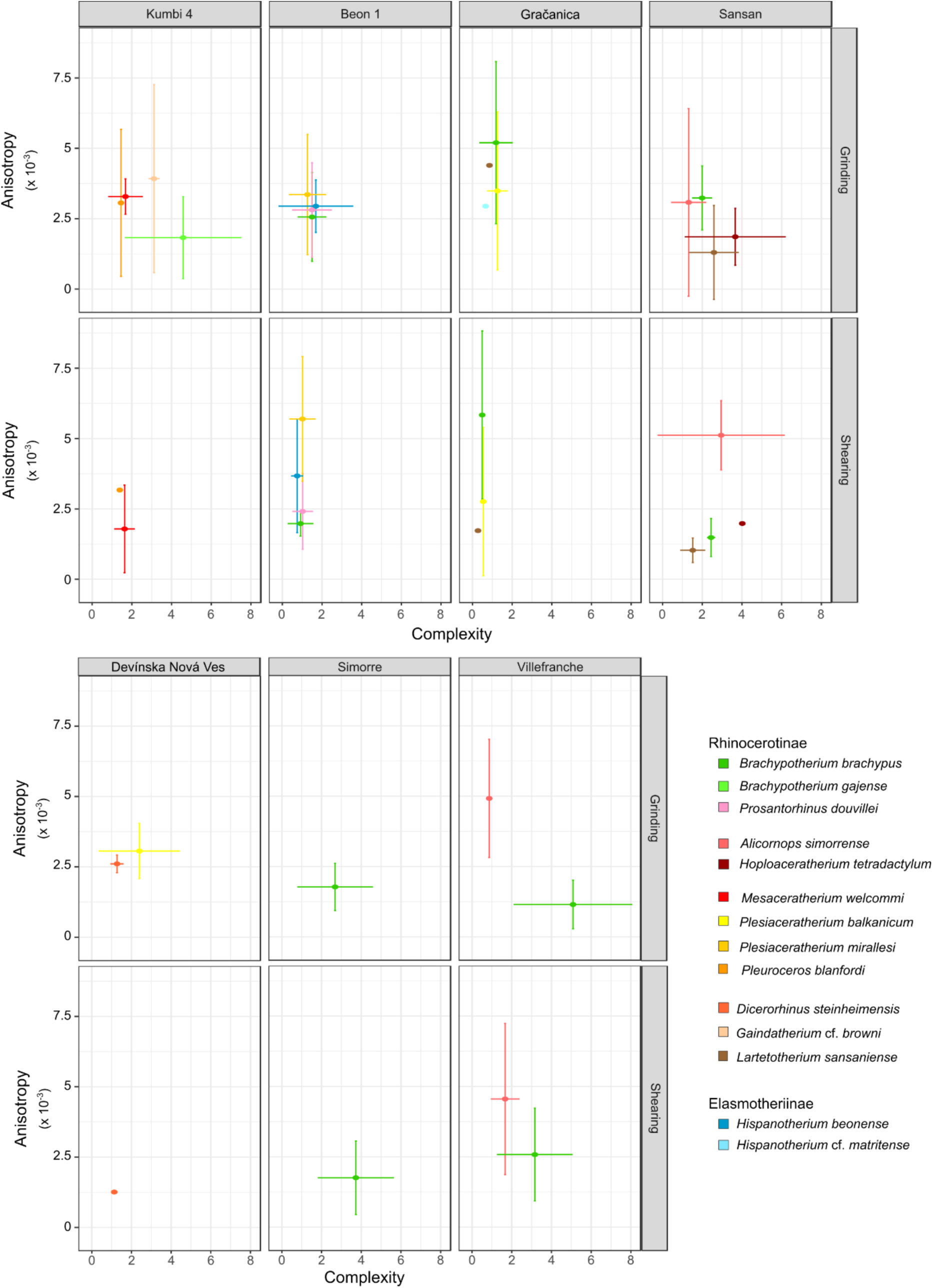
Dental microwear results of early and middle Miocene rhinocerotids plotted as mean and standard deviation of anisotropy against that of complexity by facet, locality and species. Localities organized chronologically. Béon 2 and Steinheim am Albuch not shown as not studied for microwear and only one specimen, respectively. Color code by species as indicated in the figure.

**Figure 4:**
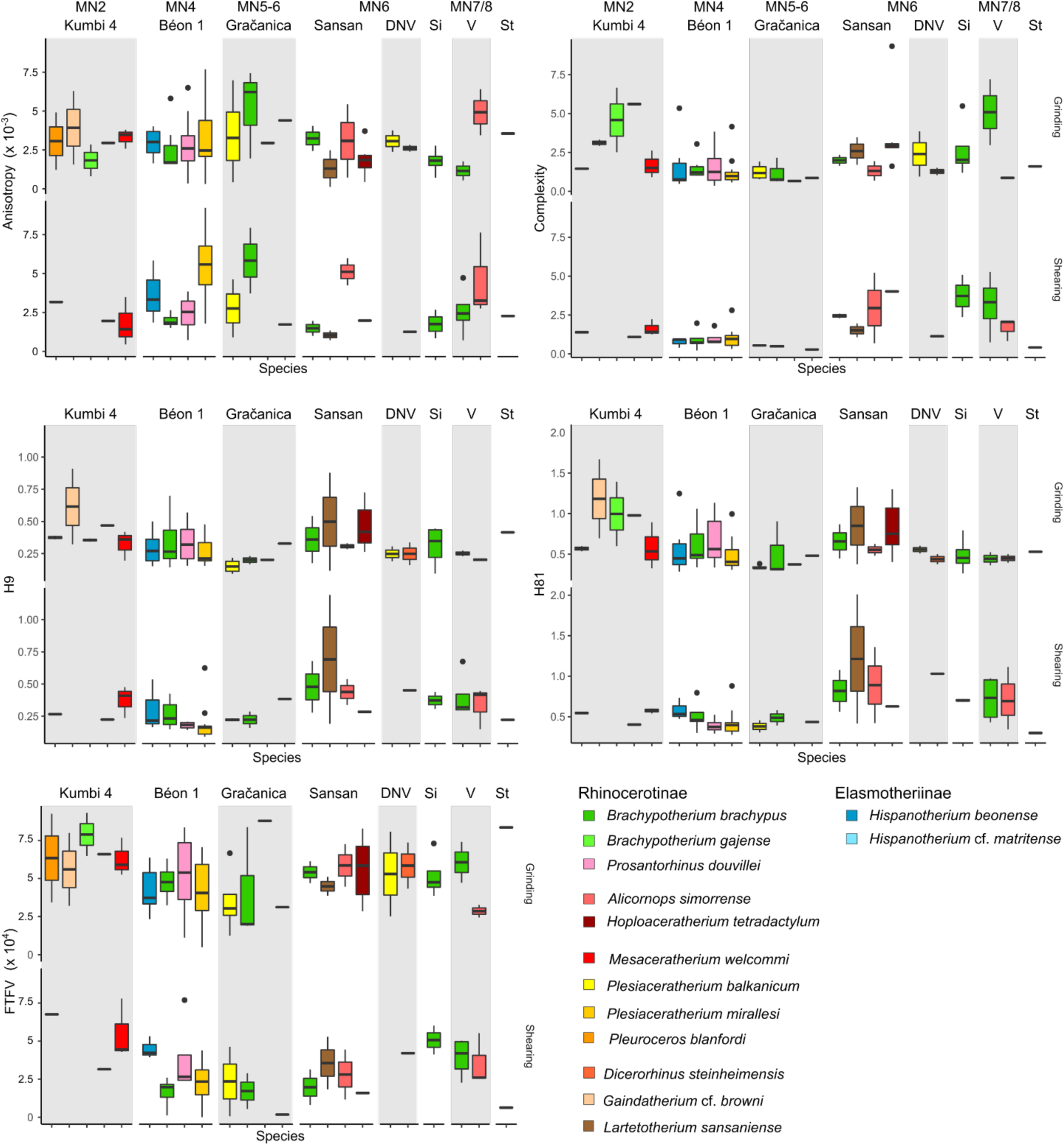
Dental microwear results of early and middle Miocene rhinocerotids plotted as boxplots of each DMTA parameter by facet and species. Time flows from left to right. DNV: Devínska Nová Ves Spalte, St: Steinheim am Albuch, Si: Simorre, V: Villefranche d’Astarac. Béon 2 not on the graph as microwear was not studied for the concerned rhinocerotids. Color code by species as indicated in the figure and consistent with Figure 3.

At **Béon 1**, the DMT patterns of the four rhinocerotids overlap contrary to that of Kumbi 4 rhinocerotids (Figure 3). The DMTA results are already detailed in Hullot et al. (2021). They suggest a mixed-feeding behavior for *H. beonense* with moderate anisotropy values (mostly < 4 x 10^-3^), variable values of complexity (low-medium), moderate-high FTfv (around 4 x 10^4^), and moderate HAsfc on both facets (HAsfc9: 0.2-0.3, HAsfc81: 0.5-0.6; Figure 4). *Plesiaceratherium mirallesi* is considered as a folivore due to low complexity (∼ 1) and HAsfc values (HAsfc9: ∼ 0.2, HAsfc81: 0.4-0.5) but relatively high anisotropy (above 5 x 10^-3^), indicating an abrasive but not diversified diet. Concerning the teleoceratines, they display similar microwear textures (Figure 3; Figure 4), although *B. brachypus* has lower values of anisotropy (< 2 x 10^-3^). This suggests that *B. brachypus* was probably a browser or a mixed-feeder, while *Pr. douvillei* was a browser favoring leaves.

At **Gračanica**, we also observe a great overlapping in the DMT patterns. The complexity is very low for all rhinocerotids studied (mostly below 1) suggesting soft food items. Anisotropy varies greatly but Hasfc81 is consistently low (< 0.5) for all species (Figure 4). This points towards soft browsing or folivory for all rhinocerotids at Gračanica.

At **Sansan**, the DMT signatures of the rhinocerotids are more diversified and less overlapping, similarly to Kumbi 4 (Figure 3). *Lartetotherium sansaniense* and *H. tetradactylum* have low values of anisotropy (< 2.5 x 10^-3^) and moderate (1-2; *L. sansaniense*) to high (> 2; *H. tetradactylum*) values of complexity, recalling browsers. The high values of HAsfc (HAsfc9 > 0.4, HAsfc81 > 0.6; Figure 4) for both species are compatible with a varied browsing diet. The other two species, *B. brachypus* and *A. simorrense*, are in the range of mixed-feeders (Figure 3), and have consistent moderate to high values of HAsfc (HAsfc9: 0.3-0.5, HAsfc81: 0.5-0.9). At **Devínska Nová Ves**, our restricted sample suggests browsing habits for both species *P. balkanicum* and *D. steinheimensis*, with moderate values of both anisotropy (∼ 2.5 x 10^-3^) and complexity (mostly between 1 and 1.5). FTfv is high on both facets (> 4 x 10^4^) and HAsfc moderate (Figure 3; Figure 4).

At **Simorre**, *B. brachypus* specimens display low values of anisotropy (< 2.5 x 10^-3^ except two specimens), and high values of complexity (> 2) and FTfv (> 4 x 10^4^) on both facets. Values of HAsfc9 are high (> 0.3) on both facets, while those of HAsfc81 are moderate on the grinding facet (median = 0.45) and high on the shearing one (median = 0.7). These DMTA results suggest browsing preferences with the inclusion of hard objects, probably fruits. At **Villefranche d’Astarac**, *B. brachypus* and *A. simorrense* present well-distinguished DMT patterns (Figure 3). *Brachypotherium brachypus* has low anisotropy values (< 2.5 x 10^-3^) and high complexity ones (> 2.5) corresponding to a browsing signal, while the opposite is true for *A. simorrense*. The moderate values of HAsfc for *A. simorrense* suggest that folivory is more likely than mixed-feeding for these specimens and the corresponding individuals. Eventually the specimen of *A. simorrense* from **Steinheim am Albuch** has a moderate anisotropy (Grinding: 3.56 x 10^-3^; Shearing: 2.27 x 10^-3^), low (Shearing: 0.41) to moderate (Grinding: 1.6) complexity, a high FTfv on the grinding facet (8.35 x 10^4^) but low on the shearing one (0.63 x 10^4^), and low HAsfc on the shearing facet (HAsfc9: 0.22, HAsfc81: 0.3) but moderate-high on the grinding one (HAsfc9: 0.42, HAsfc81: 0.53; Figure 4). This pattern is consistent with browsing or mixed-feeding habits.

#### GLMM

For all response variables (epLsar, Asfc, FTfv, HAsfc9, and HAsfc81), model support increased (i.e., lower AIC) when intraspecific factors (e.g., Facet, Genus, Locality) were included. The final models contained three to seven factors, including Specimen, the random factor, by default in all models. Facet was in the final models of epLsar and FTfv, Locality and Age were found in the final models of Asfc and both HAsfc. Details and comparison of all models can be seen in supplementary S5 and S6. Differences by Locality were also observed. The rhinocerotid specimens from Béon 1 had a lower complexity than those of Kumbi 4, Sansan, Simorre, and Villefranche (df = 119, α = 0.05, absolute t-values > 1.7), while Tukey’s contrasts highlighted lower values of Asfc for the rhinocerotids at Gračanica than for those at Kumbi (p-value = 0.004), Simorre (p-value = 0.027), and Villefranche (p-value < 0.001). Moreover, Béon 1 rhinocerotids had lower HAsfc9 and HAsfc81 values than Kumbi 4 and Sansan specimens (df = 119, α = 0.05, absolute t-values > 1.7). Tukey’s contrasts also showed that Sansan rhinocerotids had higher HAsfc9 and HAsfc81 than those at Gračanica (p-value ≤ 0.001). The sampling site (tooth locus, position, side) had sometimes a confounding effect. For instance, M2 had higher epLsar values than M3 (df = 119, α = 0.05, t-value = −1.95).

When compared to the extant dataset (see supplementary S8 for all details), we noticed that all fossil species had lower anisotropy values than the extant grazer *Ceratotherium simum* (white rhinoceros) and the folivore *Dicerorhinus sumatrensis* (Sumatran rhinoceros), although the classic t-value threshold was not reached for a few species (*P. blanfordi*, *G.* cf. *browni*, *B. gajense* [only regarding *C. simum*], *P. mirallesi*, and *A. simorrense*; α = 0.95, |t-values| ≤ 1.7). On another hand, *P. mirallesi* displayed higher values of anisotropy than the extant browsers *Diceros bicornis* (black rhinoceros; t-value = 1.93) and *Rhinoceros sondaicus* (Javan rhinoceros; t-value = 2.66). Regarding complexity, *C. simum* and *D. sumatrensis* had lower values than *B. gajense* and *H. tetradactylum*, while the extant browsers had higher values than *P. balkanicum*, *P. douvillei*, *P. mirallesi*, and *H. beonense* (α = 0.95, |t-values| > 1.7). All other DMTA parameters showed less differences between the extant and fossil datasets: *C. simum* and *R. sondaicus* had higher FTfv, HAsfc9, and HAsfc81 than *B. brachypus*, *P. balkanicum*, and *P. mirallesi* (α = 0.95, |t-values| > 1.7).

### Hypoplasia

The overall prevalence of hypoplasia on rhinocerotid teeth from the early and middle Miocene localities studied is high, with 302 teeth affected out of 1401, corresponding to over 20 % (21.56 %) all localities, teeth and species merged. There are, however, marked discrepancies between species, localities, and tooth loci (Figure 5). The most affected genera were *Plesiaceratherium* (104/357; 29.13 %), *Prosantorhinus* (97/370; 26.22 %), and *Brachypotherium* (46/178; 25.84 %), but this resulted mostly from the dominance of Béon 1 specimens in our sample. *Brachypotherium brachypus* was often one of the most affected species at all sites where the species was found, except Sansan (1/13; 7.69 %), contrary to *A. simorrense* often found associated with the previous species and relatively spared by hypoplasia (maximum 4/35 = 11.43 % of teeth affected at Simorre; Figure 6).

**Figure 5:**
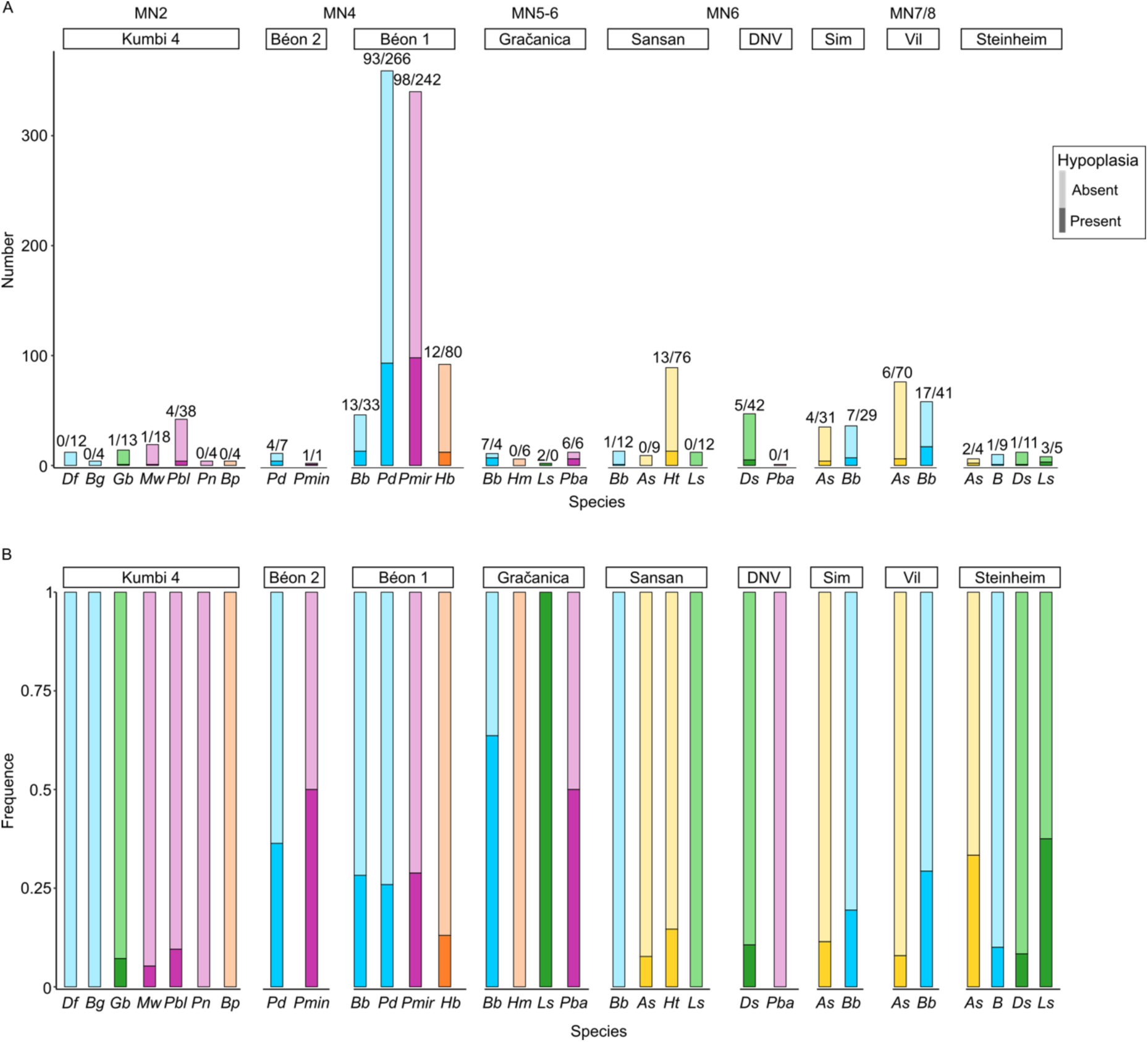
Number (A) and Frequence (B) of hypoplasia by locality and species. Numbers on barplot A indicate the number of hypoplastic teeth (dark colors) versus unaffected ones (light colors). Frequencies are calculated as the ratio of hypoplastic teeth on the total number of teeth (hypoplastic and normal). Sub-tribes colored in blue: Teleoceratina, in green: Rhinocerotina; in yellow: Aceratheriina, in pink: stem Rhinocerotinae, and in orange: Elasmotheriina Abbreviations: DNV: Devínska Nová Ves Spalte, Sim: Simorre, Vil: Villefranche d’Astarac *Df*: *Diaceratherium fatehjangense*, *Bg*: *Brachypotherium gajense*, *Bp*: *Bugtirhinus praecursor*, *Gb*: *Gaindatherium* cf. *browni*, *Mw*: *Mesaceratherium welcommi*, *Pbl*: *Pleuroceros blanfordi*, *Pn*: *Plesiaceratherium naricum*, *Pmin*: *Protaceratherium minutum*, *Pd*: *Prosantorhinus douvillei* (*P.* aff. *douvillei* at Béon 2), *Bb*: *Brachypotherium brachypus*, *Hb*: *Hispanotherium beonense*, *Pmir*: *Plesiaceratherium mirallesi*, *Hm*: *Hispanotherium* cf. *matritense*, *Ls*: *Lartetotherium sansaniense*, *Pba*: *Plesiaceratherium balkanicum*, *As*: *Alicornops simorrense*, *Ht*: *Hoploaceratherium tetradactylum*, *Ds*: *Dicerorhinus steinheimensis*, *B*: *Brachypotherium* sp.

**Figure 6:**
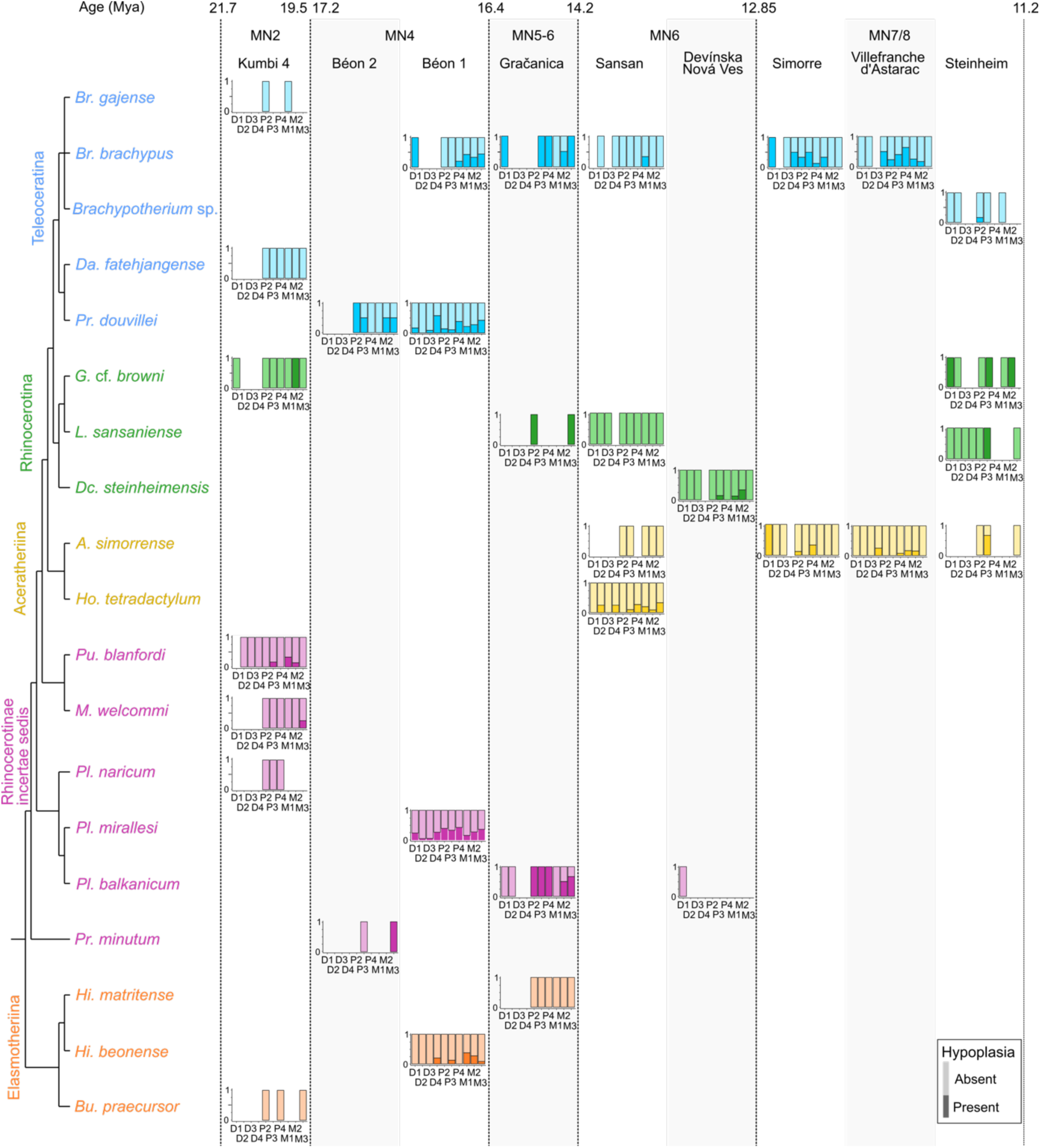
Prevalence of hypoplasia by locality, species and tooth locus plotted against phylogeny. Phylogenetic relationships follow formal parsimony analyses (Antoine, 2002; Antoine et al., 2010, 2022; Becker et al., 2013; Tissier et al., 2020). Subtribes colored in blue: Teleoceratina, in green: Rhinocerotina; in yellow: Aceratheriina, in pink: stem Rhinocerotinae, and in orange: Elasmotheriina Dark colors: hypoplastic teeth; Light colors: unaffected teeth

The prevalence was above 10 % for all localities except Kumbi 4, for which the overall prevalence is low (6/99; 6.06 %; Table 4). Hypoplasia defects are quite rare at Kumbi 4 for all species studied, and even null for the teleoceratine species (*D. fatehjangense* and *B. gajense*), *Bugtirhinus praecursor*, and *Plesiaceratherium naricum* (Figure 6). Only *Pleuroceros blanfordi* appears a little more affected (4/42; 9.52 %), totaling four of the six hypoplasias observed at the locality. Hypoplasia is also relatively limited at Sansan (14/132; 10.61 %) and Devínska Nová Ves (5/48; 10.42 %), with only *B. brachypus* and *H. tetradactylum* affected at Sansan, and *D. steinheimensis* at Devínska Nová Ves (Table 4; Figure 6). On the contrary, the rhinocerotids from Béon 1, Béon 2, and Gračanica are very affected, with more than 25 % of the teeth presenting at least one hypoplasia at Béon 1 (216/832; 25.96 %) and Béon 2 (5/18; 27.78 %), and nearly 50 % at Gračanica (15/31; 48.39 %; Table 4). At these sites, the prevalence of hypoplasia is high for all species but the elasmotheriines (Figure 6). Indeed, the elasmotheriines of all sites were relatively spared (*H. beonense* at Béon 1: 13.04 %) or even not affected by hypoplasia (*B. praecursor* at Kumbi 4 and *H.* cf. *matritense* at Gračanica).

**Table 4:**
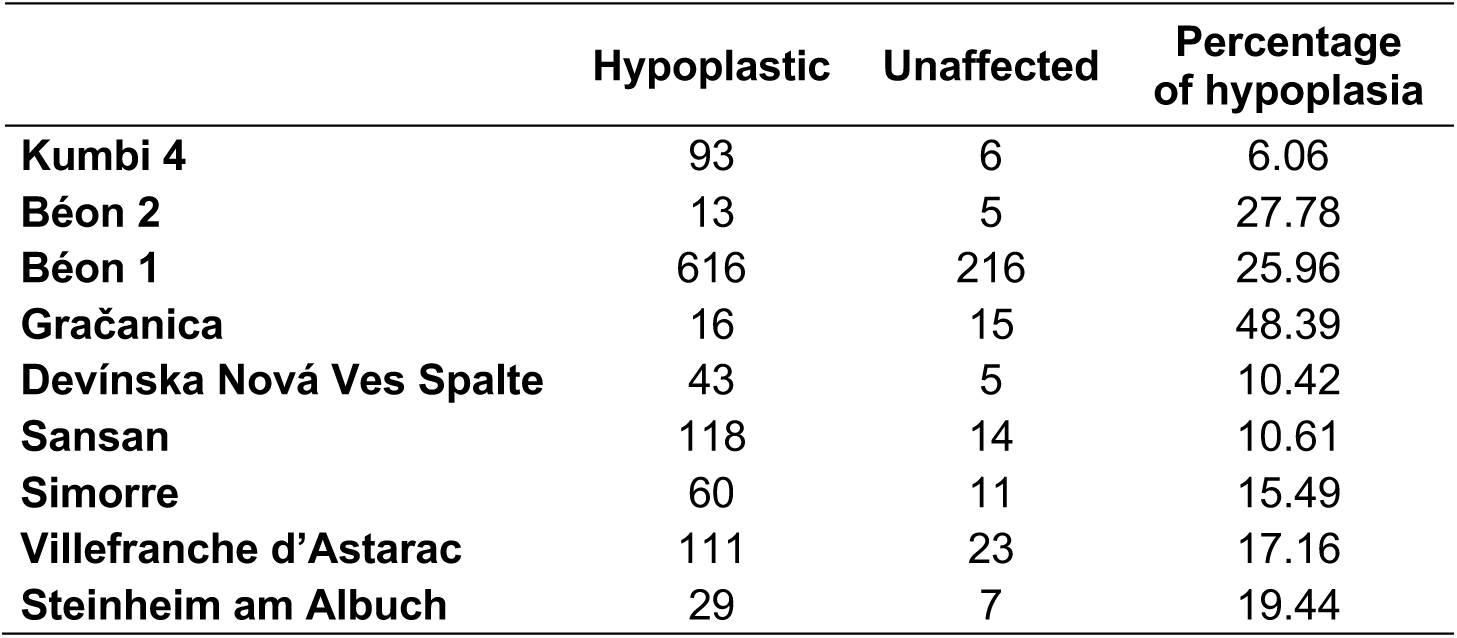
Prevalence of hypoplasia by locality (number of specimens/percentages)

Regarding the loci, milk teeth (47/294; 15.99 %) were overall less affected by hypoplasia than permanent ones (255/1107; 23.04 %; Table 5). Indeed, besides at Béon 1, very few milk molars are hypoplastic (one D1/d1 at Gračanica and Steinheim, two D1 at Simorre, two D2 and one d4 at Sansan, four D4/d4 at Villefranche d’Astarac). Upper and lower teeth were equally affected (Kruskal-Wallis, df = 1, p-value = 0.11), with respectively 19.86 % (144/725) and 23.37 % (158/676) of teeth bearing hypoplasia. The most affected locus was the fourth milk molar with 38.24 % (26/68), while the least affected were second and third milk molars with around 4 % affected (3/68 and 3/72 respectively; Table 5). Other loci particularly affected were fourth premolars (60/200; 30 %), third molars (50/188; 26.60 %), and second molars (49/202; 24.26 %; Table 5). Once again, these findings mostly result from the dominance of Béon 1 specimens in the sample, and great differences in the hypoplasia pattern are observed by locality (Figure 6). Indeed, if virtually all tooth loci are likely to be affected for Béon 1 rhinocerotids, the pattern is less varied at other localities although it seemingly diversifies with sample size (e.g., *H. tetradactylum* from Sansan and Villefranche d’Astarac). For instance, hypoplastic teeth are nearly exclusively molars at Kumbi 4 (with only one defect on a p3), and permanent teeth at Gračanica (only one defect on a D1; Figure 6).

**Table 5:** Prevalence of hypoplasia by tooth locus, regardless of provenance and taxon. Upper and lower teeth merged as they have a similar timing of development

|  | Hypoplastic | Unaffected | Percentage<br>of hypoplasia |
| --- | --- | --- | --- |
| **D1** | 71 | 15 | 17.44 |
| **D2** | 65 | 3 | 4.41 |
| **D3** | 69 | 3 | 4.17 |
| **D4** | 42 | 26 | 38.24 |
| **Total decidual teeth** | 47 | 247 | 15.99 |
| **P2** | 130 | 26 | 16.67 |
| **P3** | 138 | 33 | 19.30 |
| **P4** | 140 | 60 | 30.00 |
| **M1** | 153 | 37 | 19.47 |
| **M2** | 153 | 49 | 24.26 |
| **M3** | 138 | 50 | 26.60 |
| **Total permanent teeth** | 255 | 852 | 23.04 |

#### GLMM

For all response variables (Hypo, Defect, Multiple, Localization, and Severity), model support increased (lower AIC) when intraspecific factors (e.g., Tooth Loci, Genus, Locality) were included. When Genus was not forced into the models, the final models contained three to six factors, including Specimen, the random factor, by default in all models. Defect (converted to a factor) was in the final models of all concerned variables (Multiple, Localization, and Severity). Genus was in the final models of all variables but Localization. Position was in the final models of Hypo, Defect, and Localization, while Tooth, and Wear were in that of Hypo and Defect. Details and comparison of all models can be seen in supplementary S5 and S7.

Based on GLMMs results, we can assess the influence of Genus, Locality, and Tooth on the hypoplasia pattern. *Alicornops* was less affected than *Brachypotherium* (p-value = 0.015), while *Plesiaceratherium* was more prone to hypoplasia than *Hispanotherium* (p-value = 0.02). GLMMs revealed differences in the patterns of hypoplasia (i.e., type of defects and their frequencies) between *Brachypotherium* and the following taxa: *Dicerorhinus* (p-value = 0.0046), *Alicornops* (p-value < 0.001), and *Protaceratherium* (p-value = 0.034). Tukey’s contrasts also revealed the lowest p-values between *Alicornops* and the following taxa: *Hispanotherium* (p-value = 0.064), *Plesiaceratherium* (p-value < 0.01), *Prosantorhinus* (p-value < 0.01), *Protaceratherium* (p-value = 0.05). Eventually *Dicerorhinus* had a different hypoplasia pattern than *Plesiaceratherium* (p-value = 0.034) and *Prosantorhinus* (p-value = 0.071).

Regarding tooth loci, all teeth but fourth premolars and third molars were less affected than fourth milk molars (p-values < 0.05). The results further suggested that the most commonly affected loci were third molars, fourth premolars and fourth milk molars, while the least affected were all milk molars but the fourth. Concerning localities, Gračanica teeth were significantly more affected than Béon 1 and Sansan specimens (p-values ≈ 0.01). Middle Miocene rhinocerotids presented a hypoplasia pattern distinct from that of early Miocene ones (p-value = 0.019). Similarly to GLMMs for DMTA, we observed confounding effects. Slightly-worn teeth had less hypoplasia than average worn (p-value = 0.005) and very worn teeth (p-value = 0.026).

## Discussion

### Dietary preferences and niche partitioning of the rhinocerotids studied

The comparison of the fossil specimens DMT to those of extant ones highlighted marked differences. This suggests that the dietary spectrum of extinct rhinocerotids might have been very distinct from that observed in living species (Hullot et al., 2019). However, the microwear textures of most fossils studied are critically distinct from that of the only extant strict grazer *Ceratotherium simum*, banning such dietary preferences for the studied fossil specimens. This finding is not surprising as grasses and associated grazing ungulates expended only during latest Miocene in Eurasia (Janis, 2008). The reconstructed dietary preferences based on DMTA are presented in Table 6 by locality and by species.

**Table 6:**
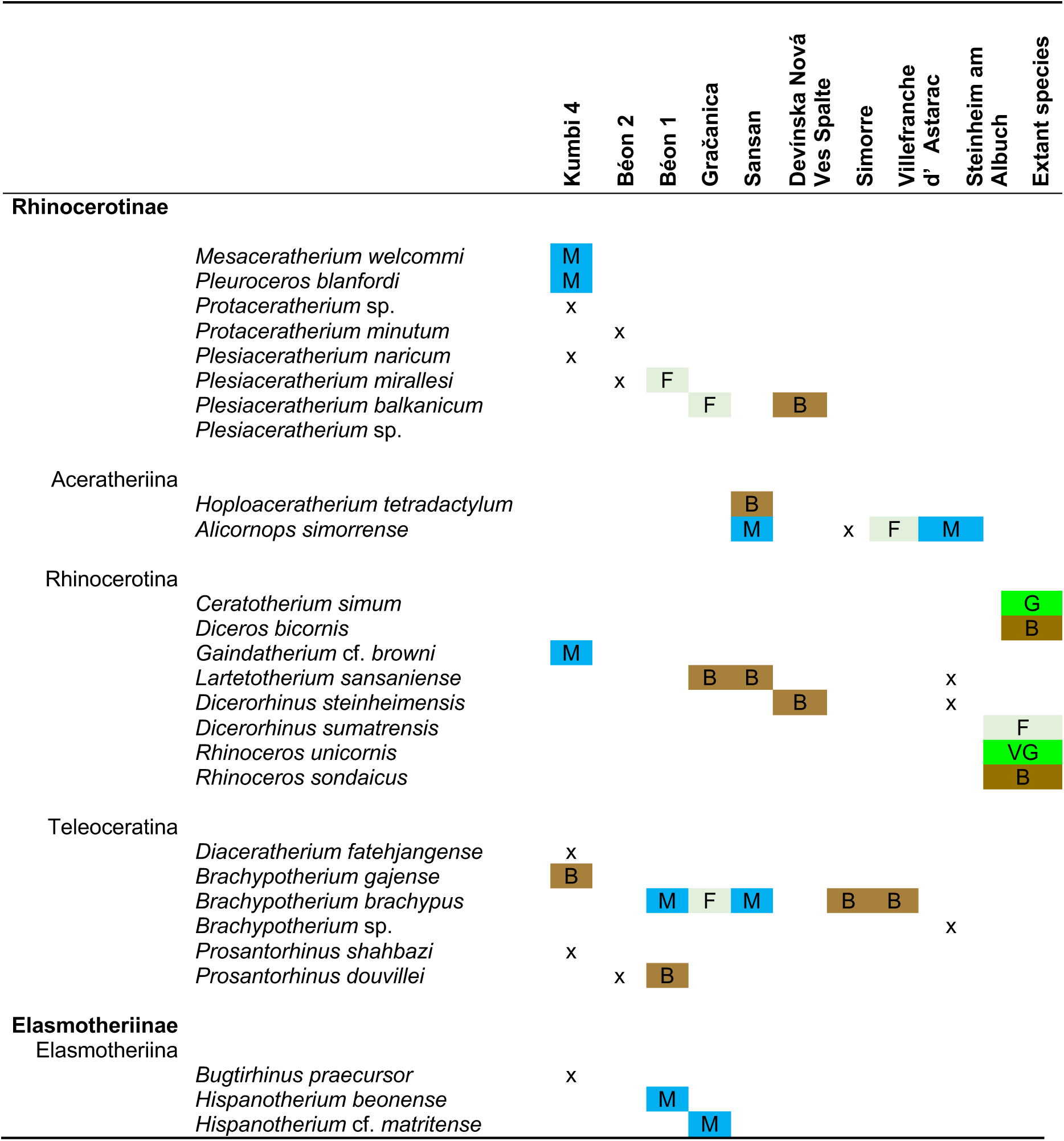
Dietary preferences inferred from textural microwear (DMTA) of the studied rhinocerotid specimens from different fossil localities of the early and middle Miocene of Eurasia. Extant species are provided as a comparison, based on the work of Hullot et al. (2019) Color code: brown/B – browser, blue/M – mixed-feeder, pale green/F – folivore, vivid green/G – grazer, vivid green/VG – variable grazer, no color/x – not studied

The DMTA results of fossil specimens on both facets (Figure 3; Figure 4) suggest clear differences in the feeding preferences of the rhinocerotid specimens studied at Kumbi 4, Sansan, and Villefranche d’Astarac, which could indicate a potential nich partitioning. Although DMT could only be explored in four out of the nine rhinocerotid species present at Kumbi 4, the patterns observed indicate clear differences in the feeding behaviors, even if leaf consumption seems to be a major component for all rhinocerotids studied but *B. gajense*. This finding is in line with the inferred lush vegetation under warm and moist climatic conditions proposed for this locality, providing abundant and diverse feeding resources for the many large herbivores present (Antoine et al., 2010, 2013; Martin et al., 2011).

The dietary preferences reconstructed for Sansan rhinocerotids suggest the co-occurrence of two browsers (*L. sansaniense* and *H. tetradactylum*, the latter including harder items in its diet) and two mixed-feeders (*A. simorrense* and *B. brachypus*), coherent with the warm (16 to 19°C mean annual temperatures) forested environment reconstructed for that locality (Costeur et al., 2012), but at odds with the recent-like Miocene coolhouse as depicted by Westerhold et al. (2020; ∼415 parts per million CO2). This discrepancy could be due to differences between local and global climatic conditions or, less probably, to a problematic dating of the locality of Sansan (Sen and Ginsburg, 2000; Maridet and Sen, 2012). Moreover, the niche partitioning was probably accentuated by different habitat preferences as suggested in the literature: *Hoploaceratherium tetradactylum* is mostly found in swamp or fluvial sediments indicating wet habitat preferences contrary to *A. simorrense*, while *B. brachypus* seems intermediate and *L. sansaniense* generalist (Heissig, 2012). Eventually, we observed obvious differences in the dietary preferences for *A. simorrense* (folivore or mixed-feeder favoring leaves) and *B. brachypus* (browser including hard objects) at Villefranche d’Astarac, where a humid forested environment is hypothesized (Bentaleb et al., 2006).

On the contrary, an overlap of microwear textures, especially for the grinding facet, is observed for the localities of Béon 1 and Gračanica. Besides diet, different habitats and feeding heights might result in niche partitioning (Hutchinson, 1959; Arsenault and Owen-Smith, 2008). Concerning Béon 1, a partial niche partitioning due to habitat differences has been hypothesized for the rhinocerotids – swamps for both teleoceratines *B. brachypus* and *P. douvillei*, open woodland for *P. mirallesi*, and savannah-like open environments for *H. beonense* (Bentaleb et al., 2006) – and subtle dietary differences are discussed in Hullot et al. (2021) in the light of the combination of molar mesowear and dental microwear texture analysis. At Gračanica, low-magnification microwear and mesowear score already revealed an overlap in the dietary preferences of *Pl. balkanicum* and *B. brachypus* as two browsing species, but the microwear pattern of the latter likely suggests dirty browsing (i.e., browsers ingesting abrasive dust, grit, or dirt with their forage; Rivals et al., 2014), a dietary category however not defined by comparative datasets on intensively studied extant species with known diet. Although microwear sampling is restricted and includes premolars for the other two rhinocerotids (*L. sansaniense* and *H.* cf. *matritense*), it points towards different mixed-feeding behaviors, most likely with a dominance of grass in the diet of *H.* cf. *matritense* (Xafis et al., 2020). The very low values of complexity for all Gračanica rhinocerotids in our sample, combined to relatively high values of anisotropy (Figure 3; Figure 4), recalls the feeding preferences and microwear textures of the extant *Dicerorhinus sumatrensis* (Sumatran rhino; Hullot et al., 2019). This could suggest an important consumption of leaves for all species, as well as a very low amount of lignified tissues that would have required more grinding to get access to cell content. Interestingly, the reconstructed environment at this locality (based on mammal assemblage and flora) is a lowland swamp surrounded by a closed canopy-like environment (Butzmann et al., 2020; Xafis et al., 2020), meaning that leaves would have been an abundant resource. Eventually, the restricted DMTA samples from Devínska Nová Ves Spalte, Steinheim am Albuch, and Simorre suggest browsing or mixed-feeding habits for all specimens studied, but did not allow to conclude on potential competition for food resources.

### Interactions with co-occurring herbivores

Besides other rhinocerotid species, the individuals studied co-occurred with many other meso- and megaherbivores (> 4 kg; bovids, cervids, giraffids, anthracotheriids, equids, chalicotheriids, and proboscideans). The co-occurrence is inferred based on the retrieving from the same stratigraphic level and/or fossil locus (Kumbi 4: Antoine et al., 2010; Gračanica: Göhlich and Mandic, 2020; Steinheim: Tütken et al., 2006) or by attested interactions between some species (Béon 1: trampling marks, Antoine pers. obs., Sansan: Aiglstorfer et al., 2019). Although co-occurrence is not necessarily a good proxy for ecological interactions (Blanchet et al., 2020), it is possible that some of these large herbivores were competing for or partitioning food resources with the rhinocerotids. Unfortunately, very little has been studied concerning the dietary preferences of the fauna at most of the studied localities with the notable exceptions of Gračanica (Göhlich and Mandic, 2020) and Sansan (Peigné and Sen, 2012).

Indeed, recent studies on dental wear (micro- and meso-wear) or stable isotopy, suggested frugivory for some associated species such as tragulids (*Dorcatherium* spp. at all localities but Simorre and Villefranche; Aiglstorfer et al., 2014; Xafis et al., 2020), the middle Miocene Moschidae (*Micromeryx* spp. found at Sansan, Simorre and Steinheim am Albuch; Aiglstorfer and Semprebon, 2019) or the chalicothere *Metaschizotherium fraasi* (found at Steinheim am Albuch; Semprebon et al., 2011). Interestingly, no rhinocerotid specimens studied here seemingly favored fruits, although they might have included some in their diet (*B. brachypus* from Simorre) or consumed some seasonally, which might not be detected by DMTA.

Similarly, the lophodont suid *Listriodon splendens*, found at Gračanica, Devínska Nová Ves, and Simorre, might have favored grasses (Van der Made, 2003; Xafis et al., 2020), a resource not dominant in the diet of the sampled rhinocerotids neither. Otherwise, the vast majority of herbivore species were probably browsers or mixed-feeders, in good agreement with the statement by Eronen and Rössner (2007) that these forms are dominant between MN4 and MN9. This is for instance the case of the associated perissodactyl species *Anisodon grande* (Chalicotheriidae), whose low-magnification microwear signal at Devínska Nová Ves suggests folivory (Semprebon et al., 2011), and *Anchitherium* spp. (Equidae) ranging from generalists to dirty browsers depending on the locality (Kaiser, 2009; Xafis et al., 2020).

Within browsers and mixed-feeders, resources’ partitioning is still possible (consumption of different plant parts or species) but might be difficult to detect in fossil communities. Moreover, other strategies can lead to niche partitioning, such as different habitat, different body mass, or different feeding height (Hutchinson, 1959; Schoener, 1974; Arsenault and Owen-Smith, 2008). Regarding body mass, most rhinocerotids studied are megaherbivores *sensu* Owen-Smith (1988; terrestrial herbivores weighting more than 1000 kg), which implies specific feeding strategies and metabolic requirements. Megaherbivores are often treated as a separate herbivore guild, mostly disturbing the feeding and abundance of mesoherbivores (4–450 kg; Fritz et al., 2002; Calandra et al., 2008; Landman et al., 2013). Within megaherbivores, proboscideans frequently co-occurred with rhinocerotids at the studied localities and were mostly browsers or mixed-feeders, placing them as direct competitors for rhinocerotids. Indeed, the mesowear and low-magnification microwear suggest that *Prodeinotherium bavaricum* and *Gomphotherium angustidens* were browsers at Gračanica (Xafis et al. 2020), while the mesowear angle categorizes *P. bavaricum* from Sansan, *D. giganteum* from Villefranche d’Astarac and *G. angustidens* from Simorre as browsers, but *G. angustidens* from Sansan and Villefranche d’Astarac as mixed-feeders (Loponen, 2020). Such overlapping in the diet of proboscideans and rhinocerotids is observed nowadays between African elephants and black rhinoceroses (Landman et al., 2013). Interestingly, this competition is detrimental to the rhinoceros, with individuals shifting towards the inclusion of more grasses in presence of elephants (on a seasonal basis). A similar shift could be hypothesized at the studied localities, notably at Gračanica, for which the low-magnification microwear of *P. bavaricum* and *L. sansaniense* would be consistent with the consumption of more grass (Xafis et al. 2020). Another possibility, as postulated by Xafis et al. (2020) for *Deinotherium* spp. and *Plesiaceratherium balkanicum* at Gračanica, would be different feeding heights between proboscideans and rhinocerotids, as the first ones were most likely feeding at the top of trees due to their larger size (some of the biggest Neogene mammals; Larramendi, 2015).

### Hypoplasia prevalence and environmental conditions

We found that the hypoplasia prevalence and pattern (i.e., tooth loci affected) were very different depending on the locality and the species concerned. Except for Kumbi 4, the prevalence was relatively high (> 10 %) at all sites of our early-middle Miocene sample. Even though nine species of rhinocerotids are found at Kumbi 4, such a low prevalence is in agreement with previous results in the region over the Cenozoic (Roohi et al., 2015), and coherent with the very favorable, low-stress context hypothesized at this locality, that is a rich vegetation under a warm and humid climate (Antoine et al., 2013).

The prevalence of hypoplasia is high at Béon 1 (>25 %) for all rhinocerotids except *H. beonense*, with molars being particularly affected with respect to other dental loci. Second and third molars are the last teeth to develop and erupt in rhinocerotids (Hitchins, 1978; Hillman-Smith et al., 1986; Böhmer et al., 2016), and stresses on these late-developed teeth have been correlated with environmental, seasonal stresses in sheep (Upex and Dobney, 2012). Although subtropical wet conditions are reconstructed at Béon 1 (just prior to the MCO), periodic droughts are also reported in the area at that time (Duranthon et al., 1999; Hullot and Antoine, 2020). Interestingly, the least affected species is the elasmotheriine *H. beonense*, an early representative of a clade adapted to relatively open and arid environments (Cerdeño and Nieto, 1995; Iñigo and Cerdeño, 1997), and which displays a mixed-feeding diet thus probably not relying on a single (or a few) specific resource(s) (Figure 4). On the contrary, both teleoceratine species, often considered swamp dwellers and hence probably depending on water availability, display a high prevalence of hypoplasia (Figure 5).

We found a very high prevalence of hypoplasia at Gračanica, with nearly 50 % of the teeth bearing at least one hypoplastic defect. The proposed age for the locality ranges between 14.8 and 13.8 Ma (Göhlich and Mandic, 2020), which is an interval of great climatic changes. Indeed, though included in the MCO, the interval from 14.7 to 14.5 Ma presents an increased seasonality in precipitations, with extended dry periods (Böhme, 2003). On the other hand, an abrupt cooling occurred between 14 and 13.5 Ma, correlating with the Mi-3 event (Zachos et al., 2001; Böhme, 2003; Holbourn et al., 2014; mMCT of Westerhold et al., 2020). Besides this challenging environmental context for the rhinocerotids, our DMTA results suggest a potential competition for food resources (Figure 3), that could have generated stressful conditions for meso- and megaherbivores.

At Sansan, the prevalence of hypoplasia is overall moderate (∼ 10 %) and defects are only found in two species out of four: *H. tetradactylum* and *B. brachypus* (only one M1). The pattern of hypoplasia for *H. tetradactylum*, with various loci affected, suggests different stresses and timing, from *in utero* (D2) to post-weaning (M3). It is quite remarkable, as the proximity of the MCO peak (Maridet and Sen, 2012) leading to seasonal warm and moist conditions (Costeur et al., 2012), would seemingly constitute relatively low stress conditions for the concerned rhinocerotids.

The prevalence of hypoplasia at Devínska Nová Ves Spalte is also moderate (5/48; 10.42 %) and restricted to *D. steinheimensis* (P3, M1, M2 only; Figure 6), although the locality dates from the mMCT. However, despite this transitional climatic system, pollen data from the Vienna Basin, to which the locality belongs, indicate that regional conditions remained tropical with few precipitation variations (Sabol and Kováč, 2006), coherent with the absence of hypoplasia on third molars, that could be correlated with seasonal stresses. The paleogeographic context seems to have played a major role, as the taxonomic differences with Sansan are partly explained by different paleoenvironments: Devínska Nová Ves Spalte was a forested area near the shoreline of the transgressive late Langhian sea (Sabol and Kováč, 2006).

The rhinocerotids from the localities of the MN7/8 (Simorre, Villefranche d’Astarac, and Steinheim am Albuch), a time of sea-level drop and comparatively dry climate (Legendre et al., 2005; Böhme et al., 2011; Heissig, 2012; Westerhold et al., 2020), presented higher prevalences but contrasted patterns depending on the species and locality (Figure 5; Figure 6). However, contrary to what we could have expected regarding the environmental conditions, the most affected loci (P2, P3, D1) document mostly early-life stresses (e.g., birth, juvenile disease), rather than environmental or seasonal stresses (Niven et al., 2004; Upex and Dobney, 2012). At Steinheim, only *L. sansaniense* has hypoplasia on other teeth than second and third premolars, suggesting mostly early-life stresses. At Simorre, more loci are affected (D1, P2-P3, P4, and M1) and the pattern is relatively similar for both co-occurring species i.e., *B. brachypus* and *A. simorrense*. Hypoplasia on D1, that develops mostly *in utero* synchronously with D4, could indicate birth-related stresses (Hillman-Smith et al., 1986; Mead, 1999; Böhmer et al., 2016). Indeed, birth is a stressful moment for most animals as it causes a temporary malnutrition and rapid environmental changes (Upex and Dobney, 2012). Similarly, the M1 starts its development relatively early, as attested by the presence of a neonatal line in some rhinocerotid teeth (Tafforeau et al., 2007), and hypoplasia on M1 could thus reveal particularly stressful conditions around birth. Eventually, the rhinocerotids from Villefranche d’Astarac document later-life stresses consistent with the dry and arid conditions mentioned above, with hypoplasia recorded from D4 to M2 (not P2-P3 for *A.simorrense*). The fourth premolars are particularly affected for *B. brachypus*, which could indicate harsh weaning or cow-calf separation conditions. Interestingly, the timing of weaning varies according to climatic conditions in proboscideans (shorter under favorable conditions), and a delayed weaning renders the individuals more vulnerable to climatic stressors (Metcalfe et al., 2010). A similar phenomenon could be hypothesized for rhinocerotids, which could result in the high prevalence of hypoplasia observed on the fourth premolars of *B. brachypus* at Villefranche.

### Paleoecological implications and changes

Several species or genera are retrieved in various localities overtime, but *B. brachypus* clearly has the longest range (from Béon 1 [MN4] to Simorre + Villefranche d’Astarac [MN7/8], with Gračanica [MN5-6] and Sansan [MN6] in the meantime). We observe a clear shift in the DMT of *B. brachypus* over time from a mixed-feeding behavior at Béon 1, Gračanica, and Sansan to a clear browsing signal with the ingestion of harder items (fruits, seeds, or even soil) at Simorre and Villefranche d’Astarac (Figure 3; Figure 4). This result could be due to a change in the regional climatic conditions, from warm and humid pre-MCO to cooler, more seasonal and arid post-MCO (Zachos et al., 2001; Böhme, 2003; Holbourn et al., 2014), perhaps leading to behavioral changes in this species (Cerdeño and Nieto, 1995), and/or to changes in local conditions. Interestingly, the δ^18^O of carbonates in the bioapatite of *brachypus* teeth at Béon 1, Sansan, and Simorre also shows marked differences, with a 2.9 ‰ decrease between MN4 and MN7/8, suggesting a decrease of the temperature of more than 4°C (Bentaleb et al., 2006). As the prevalence of hypoplasia is also very high at all localities but Sansan for this rhinocerotid, the observed dietary changes could also be due to competition with other herbivores (e.g., other rhinocerotids at Béon 1 and Gračanica, proboscideans as discussed above). Eventually, the DMT pattern at Gračanica is more singular, with high values of anisotropy (> 5 x 10^-3^). The low-magnification microwear of *B. brachypus* was also studied at Gračanica, and the authors discussed a dirty browsing behavior, consistent with the low head posture of this rhinocerotid (Becker and Tissier, 2020; Xafis et al., 2020). Such ingestion of abrasive soil particles could explain the high anisotropies observed in the studied specimens, but not the relatively low values of complexity and FTfv. Such a DMT pattern recalls that of the extant species (Scott, 2012; Ramdarshan et al., 2016), notably *D. sumatrensis* (Hullot et al., 2019), and could point towards an important consumption of leaves from low vegetation (folivory). This diet would be consistent with the abundance of trees at Gračanica (Butzmann et al., 2020; Xafis et al., 2020),

Contrastingly, the DMT of *A. simorrense* remains quite similar from Sansan (MN6) to Steinheim (MN7/8; Figure 3; Figure 4). Interestingly, there are clear differences in the hypoplasia prevalence of these two species, *B. brachypus* being one of the most affected species in our sample. Such differences in the hypoplasia prevalence could reveal the existence of a competition for food and/or water resources, that could be responsible for a shift in the diet of *B. brachypus*. The pattern of hypoplasia at Simorre (D1, P2-P3) and Villefranche d’Astarac (D4-M1, P4-M2) suggests early life stresses for both rhinocerotids (Figure 6), mostly before weaning (Mead, 1999), which does not give insight into such a competition.

Concerning other rhinocerotid species found at more than one locality (*L. sansaniense*, *D. steinheimensis*, and *Plesiaceratherium* spp.), the hypoplasia patterns seem to be different at each locality, denoting a greater effect of local conditions than species-related sensitivities. Only the *Plesiaceratherium* species from Béon 1 and Gračanica exhibit comparable patterns (Figure 6), although it could be related to the high prevalence of stresses for individuals belonging to the concerned taxa at these localities. Overall, the elasmotheriines (*Bugtirhinus, Hispanotherium*) were seemingly spared by hypoplasia. Indeed, no tooth of elasmotheres was hypoplastic at Kumbi 4 (*B. praecursor*; 0/4) and Gračanica (*H.* cf. *matritense*; 0/6). At Béon 1, for which a greater sample is available, *H. beonense* is the least affected species with only 13.04 % (12/92) of hypoplastic teeth, nearly exclusively permanent (only one hypoplasia on a D4). If this result was not surprising at Kumbi 4, where low-stress conditions were inferred and very little hypoplasia recorded for all studied species, the difference with other associated species was particularly striking at Béon 1 (prevalence above 25 % for the three other species at the site) and Gračanica (prevalence above 50 % for the three other species at the site). The microwear study of elasmotheriines is restricted in the literature, but it suggests the inclusion of a non-negligible part of browse resources in the diet, at least seasonally (Rivals et al., 2020; Xafis et al., 2020). This finding is in line with our DMTA results for *Hispanotherium* species (Béon 1 and Gračanica) suggesting mixed-feeding preferences (Figure 3; Figure 4). The increasing crown height observed in this clade over time could allow for accommodating to a greater variety of food items (Semprebon and Rivals, 2007; Damuth and Janis, 2011; Tütken et al., 2013), thus limiting nutritional stress, as observed in hipparionine equids, with respect to anchitheriine equids (MacFadden, 1992; Janis, 2008; Mihlbachler et al., 2011). The classic view of elasmotheriines as obligate open-environment rhinocerotids adapted to grazing – notably based on representatives from the arid Iberic Peninsula (Iñigo and Cerdeño, 1997) – is thus somehow challenged. This could mean that hypsodonty in this clade might counterbalance significant grit load induced by feeding low in open environments, thus reflecting more the habitat rather than the diet, an hypothesis that has already been proposed to explain hypsodonty evolution (Janis, 1988; Jardine et al., 2012; Semprebon et al., 2019). A lower prevalence of hypoplasia in elasmotheriines compared to other associated rhinocerotids has also been noted at Maragha (in the latest Miocene, though), where *Iranotherium morgani* does not present hypoplastic defects (0/16 teeth; Hullot et al., 2022). This finding could suggest an influence of diet in the stress susceptibility, with more specialized species (i.e., heavily relying on a limited number of resources) being more sensitive.

## Conclusions

In this article, we wanted to assess if the great diversity of Miocene Rhinocerotidae was associated to an ecological diversity and to explore how these paleoecological differences might reflect climate changes between the studied species, regions, and periods. The study of the paleoecology of rhinocerotids from nine localities of the early and middle Miocene of Europe and Pakistan revealed clear differences over time and space between and within species. Although the DMTA results in this sample suggested only browsers and mixed-feeders (no grazers and frugivores), we observed differences within these dietary categories that widely exceed the ecological range of extant relatives. Regarding enamel hypoplasia, which is rather prevalent in the studied sample (except in the oldest and only South Asian locality, Kumbi 4), it revealed clear disparities between localities, species, and dental loci. While the effects of global climate changes (MCO, mMCT) were often not immediately obvious, hypoplasia revealed itself as marker of more specific, local conditions at least in certain taxa (Rhinocerotina). Indeed, our sample highlighted various stress susceptibilities depending on the species and even the sub-tribe: while *B. brachypus* is highly affected by hypoplasia regardless of locality and conditions, elasmotheriines (*Bugtirhinus praecursor* at Kumbi 4, *Hispanotherium beonense* at Béon 1 and *H.* cf. *matritense* at Gračanica) are pretty spared in contrast. This could suggest an influence of phylogeny and/or diet in stress susceptibility, with more flexible species being less vulnerable to environmental stressors. The investigation of diet preferences and stress susceptibility in other taxa associated with rhinocerotids at the different localities could be of great interest to discriminate between the effects of local conditions (similar prevalences in various taxa from the same locality), phylogeny (similar prevalences in related taxa regardless of the locality) and diet in the prevalence of hypoplasias.

## Acknowledgments

We are indebted to the curators in charge of all the collections we visited and studied: U. Göhlich (NHMW, Austria), L. Costeur (NHMB, Switzerland), Y. Laurent and P. Dalous (MHNT, France). We are particularly grateful to A. Uzunidis (IPHES, Spain), C. Mallet (University of Liège, Belgium) and M.C. Mihlbachler (New York Institute of Technology, USA) for their thorough and constructive revision of an early version of the manuscript, and to A. Houssaye (Mécadev, MNHN, Paris) for her deep involvement in editing and recommending this work.

## Additional information

### Author contributions

MH collected, analyzed and interpreted the data, prepared figures, and wrote the manuscript. GM helped with the DMT analyses and the revision of the manuscript. POA conceived the study, provided taxonomic assignments and phylogenetic relationships, helped with the design of the figures and the revision of the manuscript.

### Data availability

Supplementary data are available on BioRxiv: https://www.biorxiv.org/content/10.1101/2022.05.06.490903v3.supplementary-material

## References

Aiglstorfer, M., and G. M. Semprebon. 2019. Hungry for fruit? – A case study on the ecology of middle Miocene Moschidae (Mammalia, Ruminantia). Geodiversitas, 41:385–399. doi: 10.5252/geodiversitas2019v41a10.

Aiglstorfer, M., G. E. Rössner, and M. Böhme. 2014. *Dorcatherium naui* and pecoran ruminants from the late Middle Miocene Gratkorn locality (Austria). Palaeobiodiversity and Palaeoenvironments, 94:83–123. doi: 10.1007/s12549-013-0141-9.

Antoine, P.-O. 2002. Phylogénie et évolution des Elasmotheriina (Mammalia, Rhinocerotidae). Mémoires Du Muséum National d’Histoire Naturelle, 188:5–350.

Antoine, P.-O. in press. Rhinocerotids from the Siwalik faunal sequence; In C. Badgley, D. Pilbeam, and M. Morgan (eds.), At the Foot of the Himalayas: Paleontology and Ecosystem Dynamics of the Siwalik Record of Pakistan., Johns Hopkins University Press. Baltimore.

Antoine, P.-O., and F. Duranthon. 1997. Découverte de *Protaceratherium minutum* (Mammalia, Rhinocerotidae) dans le gisement Orléanien (MN 4) de Montréal-du-Gers (Gers). Annales de Paléontologie (Vert.-Invert.), 83:201–213.

Antoine, P.-O., and J.-L. Welcomme. 2000. A New Rhinoceros From The Lower Miocene Of The Bugti Hills, Baluchistan, Pakistan: The Earliest Elasmotheriine. Palaeontology, 43:795–816. doi: 10.1111/1475-4983.00150.

Antoine, P.-O., and D. Becker. 2013. A brief review of Agenian rhinocerotids in Western Europe. Swiss Journal of Geosciences, 106:135–146. doi: 10.1007/s00015-013-0126-8.

Antoine, P.-O., F. Duranthon, and P. Tassy. 1997. L’apport des grands mammifères (Rhinocérotidés, Suoidés, Proboscidiens) à la connaissance des gisements du Miocène d’Aquitaine (France). BioChro’M97, spécial 21:581–590.

Antoine, P.-O., M. C. Reyes, N. Amano, A. P. Bautista, C.-H. Chang, J. Claude, J. De Vos, and T. Ingicco. 2022. A new rhinoceros clade from the Pleistocene of Asia sheds light on mammal dispersals to the Philippines. Zoological Journal of the Linnean Society, 194:416–430. doi: 10.1093/zoolinnean/zlab009.

Antoine, P.-O., K. F. Downing, J.-Y. Crochet, F. Duranthon, L. J. Flynn, L. Marivaux, G. Métais, A. R. Rajpar, and G. Roohi. 2010. A revision of *Aceratherium blanfordi* Lydekker, 1884 (Mammalia: Rhinocerotidae) from the Early Miocene of Pakistan: postcranials as a key. Zoological Journal of the Linnean Society, 160:139–194. doi: 10.1111/j.1096-3642.2009.00597.x.

Antoine, P.-O., G. Métais, M. Orliac, J. Crochet, L. Flynn, L. Marivaux, A. Rajpar, G. Roohi, and J. L. Welcomme. 2013. Mammalian Neogene biostratigraphy of the Sulaiman Province, Pakistan; p. 400–422. In Fossil Mammals of Asia: Neogene Biostratigraphy and Chronology. Columbia University Press doi: 10.13140/2.1.3584.5129.

Arman, S. D., T. A. A. Prowse, A. M. C. Couzens, P. S. Ungar, and G. J. Prideaux. 2019. Incorporating intraspecific variation into dental microwear texture analysis. Journal of The Royal Society Interface, 16:20180957. doi: 10.1098/rsif.2018.0957.

Arsenault, R., and N. Owen-Smith. 2008. Resource partitioning by grass height among grazing ungulates does not follow body size relation. Oikos, 117:1711–1717. doi: 10.1111/j.1600-0706.2008.16575.x.

Bates, D., M. Mächler, B. Bolker, and S. Walker. 2015. Fitting linear mixed-effects models using lme4. Journal of Statistical Software, 67:1–48. doi: 10.18637/jss.v067.i01.

Becker, D., and J. Tissier. 2020. Rhinocerotidae from the early middle Miocene locality Gračanica (Bugojno Basin, Bosnia-Herzegovina). Palaeobiodiversity and Palaeoenvironments, 100:395– 412. doi: 10.1007/s12549-018-0352-1.

Becker, D., P.-O. Antoine, and O. Maridet. 2013. A new genus of Rhinocerotidae (Mammalia, Perissodactyla) from the Oligocene of Europe. Journal of Systematic Palaeontology, 11:947– 972. doi: 10.1080/14772019.2012.699007.

Bentaleb, I., C. Langlois, C. Martin, P. Iacumin, M. Carré, P.-O. Antoine, F. Duranthon, I. Moussa, J.-J. Jaeger, and N. Barrett. 2006. Rhinocerotid tooth enamel 18O/16O variability between 23 and 12 Ma in southwestern France. Comptes Rendus Geoscience, 338:172–179. doi: 10.1016/j.crte.2005.11.007.

Berlioz, É., D. S. Kostopoulos, C. Blondel, and G. Merceron. 2018. Feeding ecology of *Eucladoceros ctenoides* as a proxy to track regional environmental variations in Europe during the early Pleistocene. Comptes Rendus Palevol, 17:320–332. doi: 10.1016/j.crpv.2017.07.002.

Blanchet, F. G., K. Cazelles, and D. Gravel. 2020. Co-occurrence is not evidence of ecological interactions. Ecology Letters, 23:1050–1063. doi: 10.1111/ele.13525.

Böhme, M. 2003. The Miocene Climatic Optimum: evidence from ectothermic vertebrates of Central Europe. Palaeogeography, Palaeoclimatology, Palaeoecology, 195:389–401. doi: 10.1016/S0031-0182(03)00367-5.

Böhme, M., A. Ilg, and M. Winklhofer. 2008. Late Miocene “washhouse” climate in Europe. Earth and Planetary Science Letters, 275:393–401. doi: 10.1016/j.epsl.2008.09.011.

Böhme, M., M. Winklhofer, and A. Ilg. 2011. Miocene precipitation in Europe: Temporal trends and spatial gradients. Palaeogeography, Palaeoclimatology, Palaeoecology, 304:212–218.

Böhmer, C., K. Heissig, and G. E. Rössner. 2016. Dental Eruption Series and Replacement Pattern in Miocene *Prosantorhinus* (Rhinocerotidae) as Revealed by Macroscopy and X-ray: Implications for Ontogeny and Mortality Profile. Journal of Mammalian Evolution, 23:265–279. doi: 10.1007/s10914-015-9313-x.

Bruch, A. A., D. Uhl, and V. Mosbrugger. 2007. Miocene climate in Europe — Patterns and evolution: A first synthesis of NECLIME. Palaeogeography, Palaeoclimatology, Palaeoecology, 253:1–7. doi: 10.1016/j.palaeo.2007.03.030.

Butler, P. M. 1952. The milk-molars of Perissodactyla, with remarks on molar occlusion. Proceedings of the Zoological Society of London, 121:777–817. doi: 10.1111/j.1096-3642.1952.tb00784.x.

Butzmann, R., U. B. Göhlich, B. Bassler, and M. Krings. 2020. Macroflora and charophyte gyrogonites from the middle Miocene Gračanica deposits in central Bosnia and Herzegovina. Palaeobiodiversity and Palaeoenvironments, 100:479–491. doi: 10.1007/s12549-018-0356-x.

Calandra, I., U. B. Göhlich, and G. Merceron. 2008. How could sympatric megaherbivores coexist? Example of niche partitioning within a proboscidean community from the Miocene of Europe. Die Naturwissenschaften, 95:831–838. doi: 10.1007/s00114-008-0391-y.

Cerdeño, E. 1998. Diversity and evolutionary trends of the Family Rhinocerotidae (Perissodactyla). Palaeogeography, Palaeoclimatology, Palaeoecology, 141:13–34. doi: 10.1016/S0031-0182(98)00003-0.

Cerdeño, E., and M. Nieto. 1995. Changes in Western European Rhinocerotidae related to climatic variations. Palaeogeography, Palaeoclimatology, Palaeoecology, 114:325–338.

Cerling, T. E., J. M. Harris, B. J. MacFadden, M. G. Leakey, J. Quade, V. Eisenmann, and J. R. Ehleringer. 1997. Global vegetation change through the Miocene/Pliocene boundary. Nature, 389:153–158. doi: 10.1038/38229.

Clementz, M. T. 2012. New insight from old bones: stable isotope analysis of fossil mammals. Journal of Mammalogy, 93:368–380. doi: 10.1644/11-MAMM-S-179.1.

Costeur, L., C. Guérin, and O. Maridet. 2012. Paléoécologie et paléoenvironnement du site miocène de Sansan; p. 661–693. *In* S. Peigné and S. Sen (eds.), Mammifères de Sansan, Mémoires du Muséum national d’Histoire naturelle. Vol. 203. Paris.

Damuth, J., and C. M. Janis. 2011. On the relationship between hypsodonty and feeding ecology in ungulate mammals, and its utility in palaeoecology. Biological Reviews, 86:733–758. doi: 10.1111/j.1469-185X.2011.00176.x.

Duranthon, F., P.-O. Antoine, C. Bulot, and J. P. Capdeville. 1999. Le Miocène inférieur et moyen continental du bassin d’Aquitaine Livret-guide de l’excursion des Journées Crouzel (10 et 11 juillet 1999). Bulletin de La Société d’histoire Naturelle de Toulouse, 135:79–91.

Eronen, J. T., and G. E. Rössner. 2007. Wetland paradise lost: Miocene community dynamics in large herbivorous mammals from the German Molasse Basin. Evolutionary Ecology Research, 9:471–494.

Fédération Dentaire Internationale. 1982. An epidemiological index of development defects of dental enamel (DDE index). International Dental Journal, 42:411–426.

Fox, J., S. Weisberg, D. Adler, D. Bates, G. Baud-Bovy, S. Ellison, D. Firth, M. Friendly, G. Gorjanc, and S. Graves. 2012. Package ‘car.’ Vienna: R Foundation for Statistical Computing,.

Fritz, H., P. Duncan, I. J. Gordon, and A. W. Illius. 2002. Megaherbivores influence trophic guilds structure in African ungulate communities. Oecologia, 131:620–625. doi: 10.1007/s00442-002-0919-3.

Giaourtsakis, I., G. Theodorou, S. Roussiakis, A. Athanassiou, and G. Iliopoulos. 2006. Late Miocene horned rhinoceroses (Rhinocerotinae, Mammlia) from Kerassia (Euboea, Greece). Neues Jahrbuch Für Geologie Und Paläontologie - Abhandlungen, 239:367–398. doi: 10.1127/njgpa/239/2006/367.

Göhlich, U. B., and O. Mandic. 2020. Introduction to the special issue “The drowning swamp of Gračanica (Bosnia-Herzegovina)—a diversity hotspot from the middle Miocene in the Bugojno Basin.” Palaeobiodiversity and Palaeoenvironments, 100:281–293. doi: 10.1007/s12549-020-00437-0.

Goodman, A. H., and J. C. Rose. 1990. Assessment of systemic physiological perturbations from dental enamel hypoplasias and associated histological structures. American Journal of Physical Anthropology, 33:59–110. doi: 10.1002/ajpa.1330330506.

Grine, F. E. 1986. Dental evidence for dietary differences in *Australopithecus* and *Paranthropus*: a quantitative analysis of permanent molar microwear. Journal of Human Evolution, 15:783–822.

Heissig, K. 2012. Les Rhinocerotidae (Perissodactyla) de Sansan; p. 317–485. *In* S. Peigné and S. Sen (eds.), Mammifères de Sansan. Vol. 203. Mémoires du Muséum national d’Histoire naturelle, Paris.

Hillman-Smith, A. K. K., N. R. Owen-Smith, J. L. Anderson, A. J. Hall-Martin, and J. P. Selaladi. 1986. Age estimation of the white rhinoceros (*Ceratotherium simum*). Journal of Zoology, 210:355– 377.

Hitchins, P. M. 1978. Age determination of the black rhinoceros (*Diceros bicornis* Linn.) in Zululand. South African Journal of Wildlife Research, 8:71–80.

Hoffman, J. M., D. Fraser, and M. T. Clementz. 2015. Controlled feeding trials with ungulates: a new application of in vivo dental molding to assess the abrasive factors of microwear. The Journal of Experimental Biology, 218:1538–1547. doi: 10.1242/jeb.118406.

Holbourn, A., W. Kuhnt, M. Lyle, L. Schneider, O. Romero, and N. Andersen. 2014. Middle Miocene climate cooling linked to intensification of eastern equatorial Pacific upwelling. Geology, 42:19–22. doi: 10.1130/G34890.1.

Hullot, M., and P.-O. Antoine. 2020. Mortality curves and population structures of late early Miocene Rhinocerotidae (Mammalia, Perissodactyla) remains from the Béon 1 locality of Montréal-du-Gers, France. Palaeogeography, Palaeoclimatology, Palaeoecology, 558:109938. doi: 10.1016/j.palaeo.2020.109938.

Hullot, M., P.-O. Antoine, M. Ballatore, and G. Merceron. 2019. Dental microwear textures and dietary preferences of extant rhinoceroses (Perissodactyla, Mammalia). Mammal Research, 64:397– 409. doi: 10.1007/s13364-019-00427-4.

Hullot, M., Y. Laurent, G. Merceron, and P.-O. Antoine. 2021. Paleoecology of the Rhinocerotidae (Mammalia, Perissodactyla) from Béon 1, Montréal-du-Gers (late early Miocene, SW France): Insights from dental microwear texture analysis, mesowear, and enamel hypoplasia. Palaeontologia Electronica, 24:1–26. doi: 10.26879/1163.

Hullot, M., P.-O. Antoine, N. Spassov, G. D. Koufos, and G. Merceron. 2022. Late Miocene rhinocerotids from the Balkan-Iranian province: ecological insights from dental microwear textures and enamel hypoplasia. Historical Biology, NA:1–18. doi: 10.1080/08912963.2022.2095910.

Hutchinson, G. E. 1959. Homage to Santa Rosalia or why are there so many kinds of animals? The American Naturalist, 93:145–159.

Iñigo, C., and E. Cerdeño. 1997. The *Hispanotherium matritense* (Rhinocerotidae) from Córcoles (Guadalajara, Spain): Its contribution to the systematics of the Miocene Iranotheriina. Geobios, 30:243–266. doi: 10.1016/S0016-6995(97)80232-X.

Janis, C. 2008. An Evolutionary History of Browsing and Grazing Ungulates; p. 21–45. In I. J. Gordon and H. H. T. Prins (eds.), The Ecology of Browsing and Grazing. Ecological Studies Springer, Berlin, Heidelberg doi: 10.1007/978-3-540-72422-3_2.

Janis, C. M. 1988. An estimation of tooth volume and hypsodonty indices in ungulate mammals, and the correlation of these factors with dietary preferences. Memoires Du Museum National d’Histoire Naturelle, serie C, 53:367–387.

Jardine, P. E., C. M. Janis, S. Sahney, and M. J. Benton. 2012. Grit not grass: Concordant patterns of early origin of hypsodonty in Great Plains ungulates and Glires. Palaeogeography, Palaeoclimatology, Palaeoecology, 365–366:1–10. doi: 10.1016/j.palaeo.2012.09.001.

Jones, D. B., and L. R. G. DeSantis. 2017. Dietary ecology of ungulates from the La Brea tar pits in southern California: A multi-proxy approach. Palaeogeography, Palaeoclimatology, Palaeoecology, 466:110–127. doi: 10.1016/j.palaeo.2016.11.019.

Kaiser, T. M. 2009. *Anchitherium aurelianense* (Equidae, Mammalia): a brachydont “dirty browser” in the community of herbivorous large mammals from Sandelzhausen (Miocene, Germany). Paläontologische Zeitschrift, 83:131. doi: 10.1007/s12542-009-0002-z.

Landman, M., D. S. Schoeman, and G. I. H. Kerley. 2013. Shift in Black Rhinoceros Diet in the Presence of Elephant: Evidence for Competition? PLOS ONE, 8:e69771. doi: 10.1371/journal.pone.0069771.

Larramendi, A. 2015. Shoulder height, body mass, and shape of proboscideans. Acta Palaeontologica Polonica, 61:537–574. doi: 10.4202/app.00136.2014.

Legendre, S., S. Montuire, O. Maridet, and G. Escarguel. 2005. Rodents and climate: A new model for estimating past temperatures. Earth and Planetary Science Letters, 235:408–420. doi: 10.1016/j.epsl.2005.04.018.

Loponen, L. 2020. Diets of Miocene proboscideans from Eurasia, and their connection to environments and vegetation. Master Thesis, University of Helsinki, Finland, 54 p.

Louail, M., S. Ferchaud, A. Souron, A. E. C. Walker, and G. Merceron. 2021. Dental microwear textures differ in pigs with overall similar diets but fed with different seeds. Palaeogeography, Palaeoclimatology, Palaeoecology, 572:110415. doi: 10.1016/j.palaeo.2021.110415.

MacFadden, B. 1992. Fossil Horses: Systematics, Paleobiology, and Evolution of the Family Equidae, Cambridge University Press. New York, 369 p.

Maridet, O., and S. Sen. 2012. Les Cricetidae (Rodentia) de Sansan; p. 29–65. *In* Mammifères de Sansan. Vol. 203. Mémoires du Muséum Paris.

Maridet, O., G. Escarguel, L. Costeur, P. Mein, M. Hugueney, and S. Legendre. 2007. Small mammal (rodents and lagomorphs) European biogeography from the Late Oligocene to the mid Pliocene. Global Ecology and Biogeography, 16:529–544. doi: 10.1111/j.1466-8238.2006.00306.x.

Martin, C., I. Bentaleb, and P.-O. Antoine. 2011. Pakistan mammal tooth stable isotopes show paleoclimatic and paleoenvironmental changes since the early Oligocene. Palaeogeography Palaeoclimatology Palaeoecology, 311:19–29. doi: 10.1016/j.palaeo.2011.07.010.

Mead, A. J. 1999. Enamel hypoplasia in Miocene rhinoceroses (*Teleoceras*) from Nebraska: evidence of severe physiological stress. Journal of Vertebrate Paleontology, 19:391–397.

Merceron, G., A. Kallend, A. Francisco, M. Louail, F. Martin, C.-A. Plastiras, G. Thiery, and J.-R. Boisserie. 2021. Further away with dental microwear analysis: Food resource partitioning among Plio-Pleistocene monkeys from the Shungura Formation, Ethiopia. Palaeogeography, Palaeoclimatology, Palaeoecology, 572:110414. doi: 10.1016/j.palaeo.2021.110414.

Merceron, G., A. Ramdarshan, C. Blondel, J.-R. Boisserie, N. Brunetiere, A. Francisco, D. Gautier, X. Milhet, A. Novello, and D. Pret. 2016. Untangling the environmental from the dietary: dust does not matter. Proc. R. Soc. B, 283 doi: 10.1098/rspb.2016.1032.

Metcalfe, J. Z., F. J. Longstaffe, and G. D. Zazula. 2010. Nursing, weaning, and tooth development in woolly mammoths from Old Crow, Yukon, Canada: Implications for Pleistocene extinctions. Palaeogeography, Palaeoclimatology, Palaeoecology, 298:257–270. doi: 10.1016/j.palaeo.2010.09.032.

Mihlbachler, M. C., F. Rivals, N. Solounias, and G. M. Semprebon. 2011. Dietary change and evolution of horses in North America. Science, 331:1178–1181. doi: 10.1126/science.1196166.

Mihlbachler, M. C., D. Campbell, M. Ayoub, C. Chen, and I. Ghani. 2016. Comparative dental microwear of ruminant and perissodactyl molars: Implications for paleodietary analysis of rare and extinct ungulate clades. Paleobiology, 42:98–116. doi: 10.1017/pab.2015.33.

Mihlbachler, M. C., D. Campbell, C. Chen, M. Ayoub, and P. Kaur. 2018. Microwear–mesowear congruence and mortality bias in rhinoceros mass-death assemblages. Paleobiology, 44:131– 154. doi: 10.1017/pab.2017.13.

Niven, L. B., C. P. Egeland, and L. C. Todd. 2004. An inter-site comparison of enamel hypoplasia in bison: implications for paleoecology and modeling Late Plains Archaic subsistence. Journal of Archaeological Science, 31:1783–1794. doi: 10.1016/j.jas.2004.06.001.

Owen-Smith, N. R. 1988. Megaherbivores: The Influence of Very Large Body Size on Ecology. Cambridge University Press, 392 p.

Peigné, S., and S. Sen. 2012. Mammifères de Sansan, Mémoires du Muséum national d’Histoire naturelle. Muséum national d’Histoire naturelle, Paris, 709 p.

Prothero, D. R. 2005. The Evolution of North American Rhinoceroses. Cambridge University Press, 232 p.

Prothero, D. R., C. Guérin, and E. Manning. 1989. The history of the Rhinocerotoidea; p. 322–340. In D. R. Prothero and R. M. Schoch (eds.), The Evolution of Perissodactyls. Oxford University Press, New York.

Ramdarshan, A., C. Blondel, D. Gautier, J. Surault, and G. Merceron. 2017. Overcoming sampling issues in dental tribology: Insights from an experimentation on sheep. Palaeontologia Electronica, 20:1–19.

Ramdarshan, A., C. Blondel, N. Brunetière, A. Francisco, D. Gautier, J. Surault, and G. Merceron. 2016. Seeds, browse, and tooth wear: a sheep perspective. Ecology and Evolution, 6:5559– 5569. doi: 10.1002/ece3.2241.

Rivals, F., G. Semprebon, and A. Lister. 2012. An examination of dietary diversity patterns in Pleistocene proboscideans (*Mammuthus*, *Palaeoloxodon*, and *Mammut*) from Europe and North America as revealed by dental microwear. Quaternary International, 255:188–195. doi: 10.1016/j.quaint.2011.05.036.

Rivals, F., S. Takatsuki, R. M. Albert, and L. Macià. 2014. Bamboo feeding and tooth wear of three sika deer (*Cervus nippon*) populations from northern Japan. Journal of Mammalogy, 95:1043– 1053. doi: 10.1644/14-MAMM-A-097.

Rivals, F., N. E. Prilepskaya, R. I. Belyaev, and E. M. Pervushov. 2020. Dramatic change in the diet of a late Pleistocene *Elasmotherium* population during its last days of life: Implications for its catastrophic mortality in the Saratov region of Russia. Palaeogeography, Palaeoclimatology, Palaeoecology, 556:109898. doi: 10.1016/j.palaeo.2020.109898.

Roohi, G., S. M. Raza, A. M. Khan, R. M. Ahmad, and M. Akhtar. 2015. Enamel Hypoplasia in Siwalik Rhinocerotids and its Correlation with Neogene Climate. Pakistan Journal of Zoology, 47:1433–1443.

Sabol, M., and M. Kováč. 2006. Badenian palaeoenvironment, faunal succession and biostratigraphy: a case study from northern Vienna Basin, Devínska Nová Ves-Bonanza site (Western Carpathians, Slovakia). Beiträge Zur Paläontologie, 30:415–425.

Scott, J. R. 2012. Dental microwear texture analysis of extant African Bovidae. Mammalia, 76:157– 174. doi: 10.1515/mammalia-2011-0083.

Scott, R. S., P. S. Ungar, T. S. Bergstrom, C. A. Brown, F. E. Grine, M. F. Teaford, and A. Walker. 2005. Dental microwear texture analysis shows within-species diet variability in fossil hominins. Nature, 436:693–695. doi: 10.1038/nature03822.

Scott, R. S., P. S. Ungar, T. S. Bergstrom, C. A. Brown, B. E. Childs, M. F. Teaford, and A. Walker. 2006. Dental microwear texture analysis: technical considerations. Journal of Human Evolution, 51:339–349. doi: 10.1016/j.jhevol.2006.04.006.

Semprebon, G. M., and F. Rivals. 2007. Was grass more prevalent in the pronghorn past? An assessment of the dietary adaptations of Miocene to Recent Antilocapridae (Mammalia: Artiodactyla). Palaeogeography, Palaeoclimatology, Palaeoecology, 253:332–347. doi: 10.1016/j.palaeo.2007.06.006.

Semprebon, G. M., P. J. Sise, and M. C. Coombs. 2011. Potential bark and fruit browsing as revealed by stereomicrowear analysis of the peculiar clawed herbivores known as chalicotheres (Perissodactyla, Chalicotherioidea). Journal of Mammalian Evolution, 18:33–55. doi: 10.1007/s10914-010-9149-3.

Semprebon, G. M., F. Rivals, and C. M. Janis. 2019. The Role of Grass vs. Exogenous Abrasives in the Paleodietary Patterns of North American Ungulates. Frontiers in Ecology and Evolution, 7. doi: 10.3389/fevo.2019.00065.

Sen, S., and L. Ginsburg. 2000. La magnétostratigraphie du site de Sansan. Memoires Du Museum National d’Histoire Natural de Paris, 183:69–81.

Stefaniak, K., R. Stachowicz-Rybka, R. K. Borówka, A. Hrynowiecka, A. Sobczyk, M. M. Hoyo, A. Kotowski, D. Nowakowski, M. T. Krajcarz, E. M. E. Billia, D. Persico, E. M. Burkanova, S. V. Leschinskiy, E. van Asperen, U. Ratajczak, A. V. Shpansky, M. Lempart, B. Wach, M. Niska, J. van der Made, K. Stachowicz, J. Lenarczyk, J. Piątek, and O. Kovalchuk. 2020. Browsers, grazers or mix-feeders? Study of the diet of extinct Pleistocene Eurasian forest rhinoceros *Stephanorhinus kirchbergensis* (Jäger, 1839) and woolly rhinoceros *Coelodonta antiquitatis* (Blumenbach, 1799). Quaternary International, 605–606:192-212. doi: 10.1016/j.quaint.2020.08.039.

Tafforeau, P., I. Bentaleb, J.-J. Jaeger, and C. Martin. 2007. Nature of laminations and mineralization in rhinoceros enamel using histology and X-ray synchrotron microtomography: potential implications for palaeoenvironmental isotopic studies. Palaeogeography, Palaeoclimatology, Palaeoecology, 246:206–227. doi: 10.1016/j.palaeo.2006.10.001.

Tissier, J., P.-O. Antoine, and D. Becker. 2020. New material of *Epiaceratherium* and a new species of *Mesaceratherium* clear up the phylogeny of early Rhinocerotidae (Perissodactyla). Royal Society Open Science, 7:200633. doi: 10.1098/rsos.200633.

Tütken, T., T. W. Vennemann, H. Janz, and E. P. J. Heizmann. 2006. Palaeoenvironment and palaeoclimate of the Middle Miocene lake in the Steinheim basin, SW Germany: A reconstruction from C, O, and Sr isotopes of fossil remains. Palaeogeography, Palaeoclimatology, Palaeoecology, 241:457–491. doi: 10.1016/j.palaeo.2006.04.007.

Tütken, T., T. M. Kaiser, T. Vennemann, and G. Merceron. 2013. Opportunistic Feeding Strategy for the Earliest Old World Hypsodont Equids: Evidence from Stable Isotope and Dental Wear Proxies. PLOS ONE, 8:e74463. doi: 10.1371/journal.pone.0074463.

Upex, B., and K. Dobney. 2012. Dental enamel hypoplasia as indicators of seasonal environmental and physiological impacts in modern sheep populations: a model for interpreting the zooarchaeological record. Journal of Zoology, 287:259–268. doi: 10.1111/j.1469-7998.2012.00912.x.

Van der Made, J. 2003. Suoidea (Artiodactyla); p. 308–327. In Geology and paleontology of the Miocene Sinap Formation, New York (Columbia University Press).

Venables, W. N., and B. D. Ripley. 2002. Modern Applied Statistics with S, Springer. New York, 498 p.

Wang, B., and R. Secord. 2019. Paleoecology of *Aphelops* and *Teleoceras* (Rhinocerotidae) through an interval of changing climate and vegetation in the Neogene of the Great Plains, central United States. Palaeogeography, Palaeoclimatology, Palaeoecology, 109411. doi: 10.1016/j.palaeo.2019.109411.

Wasserstein, R. L., and N. A. Lazar. 2016. The ASA Statement on p-Values: Context, Process, and Purpose. The American Statistician, 70:129–133. doi: 10.1080/00031305.2016.1154108.

Wasserstein, R. L., A. L. Schirm, and N. A. Lazar. 2019. Moving to a World Beyond “p < 0.05.” The American Statistician, 73:1–19. doi: 10.1080/00031305.2019.1583913.

Westerhold, T., N. Marwan, A. J. Drury, D. Liebrand, C. Agnini, E. Anagnostou, J. S. K. Barnet, S. M. Bohaty, D. D. Vleeschouwer, F. Florindo, T. Frederichs, D. A. Hodell, A. E. Holbourn, D. Kroon, V. Lauretano, K. Littler, L. J. Lourens, M. Lyle, H. Pälike, U. Röhl, J. Tian, R. H. Wilkens, P. A. Wilson, and J. C. Zachos. 2020. An astronomically dated record of Earth’s climate and its predictability over the last 66 million years. Science, 369:1383–1387. doi: 10.1126/science.aba6853.

Wickham, H. 2007. Reshaping data with the reshape package. Journal of Statistical Software, 21:1– 20. doi: 10.18637/jss.v021.i12.

Wickham, H. 2011. ggplot2. Wiley Interdisciplinary Reviews: Computational Statistics, 3:180–185. doi: 10.1002/wics.147.

Wickham, H., R. François, L. Henry, and K. Müller. 2019. dplyr: A Grammar of Data Manipulation. R Package Version 0.8.3, 13:2020.

Winkler, D. E., E. Schulz-Kornas, T. M. Kaiser, D. Codron, J. Leichliter, J. Hummel, L. F. Martin, M. Clauss, and T. Tütken. 2020. The turnover of dental microwear texture: Testing the” last supper” effect in small mammals in a controlled feeding experiment. Palaeogeography, Palaeoclimatology, Palaeoecology, 557:109930. doi: 10.1016/j.palaeo.2020.109930.

Xafis, A., J. Saarinen, K. Bastl, D. Nagel, and F. Grímsson. 2020. Palaeodietary traits of large mammals from the middle Miocene of Gračanica (Bugojno Basin, Bosnia-Herzegovina). Palaeobiodiversity and Palaeoenvironments, 100:457–477. doi: 10.1007/s12549-020-00435-2.

Zachos, J., M. Pagani, L. Sloan, E. Thomas, and K. Billups. 2001. Trends, rhythms, and aberrations in global climate 65 Ma to present. Science, 292:686–693. doi: 10.1126/science.1059412.

