## Supplementary material for "Spatio-temporal diversity of dietary preferences and stress sensibilities of early and middle Miocene Rhinocerotidae from Eurasia: Impact of climate changes": S2

**Supplementary 2 – Material and methods**

Details on the localities are given hereunder, following a chronological order:

**Kumbi 4** is a very rich fossil locality situated in the Bugti Hills (upper member of Chitarwata Formation; Pakistan). It has been dated to the earliest Miocene, probably to the MN2 or early MN3 (Welcomme et al., 1997; Métais et al., 2009; Antoine et al., 2010, 2013, in press.). This locality bears the maximum number of co-occurring rhinocerotid species at the world scale, with specimens belonging to nine distinct species found in association (Table 1): the early elasmotheriine *Bugtirhinus praecursor*, the hornless rhinocerotines *Protaceratherium sp*. and *Plesiaceratherium naricum*, the basal rhinocerotines *Pleuroceros blanfordi* and *Mesaceratherium welcommi*, the teleoceratines *Brachypotherium gajense*, *Diaceratherium fatehjangense*, and *Prosantorhinus shahbazi*, and the rhinocerotine *Gaindatherium* cf*. browni* (Antoine and Welcomme, 2000; Antoine et al., 2010, 2013). No floral nor palynological data are available for this site, but the abundance of large herbivore species suggest a rich and varied vegetation, coherent with the favorable climatic conditions of the early Miocene (Métais et al., 2009; Antoine et al., 2010, 2013). The values of the δ^18^O of mammal enamel suggest wet conditions and a tropical forest (Martin et al., 2011).

Originally referred to as Montréal-du-Gers (Occitanie, France), this locality has been split into two different sites – **Béon 1** and **Béon 2** – due to substantial dating differences. **Béon 2** is dated to the base of the MN4 and coeval to Artenay, while **Béon 1** is younger, by mid-MN4 times (17 Mya; Antoine et al., 2018). The faunal assemblages at the two sites are also quite different, with Béon 2 having yielded a less abundant and diverse fauna, including only three rhinocerotid species: *Protaceratherium minutum*, *Prosantorhinus* aff. *douvillei,* and *Plesiaceratherium mirallesi* (Antoine and Duranthon, 1997; Antoine et al., 2018). On the other hand, more than 60 vertebrate species were found at Béon 1, among which five rhinocerotids: the teleoceratines *Brachypotherium brachypus* and *Prosantorhinus douvillei*, the basal rhinocerotines *Plesiaceratherium mirallesi* and *Plesiaceratherium* sp*.*, and the early elasmotheriine *Hispanotherium beonense* (Crouzel et al., 1988; Antoine, 1997, 2002; Antoine and Duranthon, 1997; Rage and Bailón, 2005; Orliac et al., 2006). At Béon 1, an oxbow lake or a swamp surrounded by a wooded savannah, under hot and moist climatic conditions, is proposed (Duranthon et al., 1999; Rage and Bailón, 2005).

**Gračanica** is a locality from the Bugojno Basin in Bosnia-Herzegovina. It has been dated to the early middle Miocene between 14.8 and 13.7 Mya, probably correlating with MN5 or early MN6 (Göhlich and Mandic, 2020). Four species of rhinocerotids were recognized at the locality (Table 1): *Lartetotherium sansaniense*, *Brachypotherium brachypus*, *Plesiaceratherium balkanicum* (type locality), and *Hispanotherium* cf. *matritense* (Becker and Tissier, 2020). However, if the Gračanica fauna is species-rich (29 mammalian taxa), it is poor in terms of individuals (Göhlich and Mandic, 2020). The environmental reconstruction is quite similar to that of Béon 1, as a swamp or lake with changing levels, surrounded by mosaic habitats (Becker and Tissier, 2020; Göhlich and Mandic, 2020; Xafis et al., 2020).

**Sansan** is the reference locality for the MN6. It is one of the richest European Miocene localities situated in the Gers (Occitanie, France) and has a controversial datation (Costeur et al., 2012; Peigné and Sen, 2012), although 14.1 Mya is considered most likely at the moment (Aiglstorfer et al., 2019). It has yielded 85 mammalian species (9 orders, 30 families, 75 genera), including four distinct rhincerotid species: *Brachypotherium brachypus*, *Lartetotherium sansaniense*, *Hoploaceratherium tetradactylum*, and *Alicornops simorrense* (Antoine et al., 1997; Heissig, 2012). Faunal and floral components suggest a subtropical to tropical forest with open zones near a water body, under warm but probably seasonal climatic conditions (Costeur et al., 2012).

**Devínska Nová Ves Spalte** refers to several middle Miocene localities from Bratislava, Slovakia, but with a good spatio-temporal affinity. The localities are correlated with the MN6 (Sabol and Kováč, 2006; Fejfar and Sabol, 2009). The mammal assemblage is relatively distinct from that of Sansan, probably linked to a regional marine transgression and associated environmental changes (Sabol and Kováč, 2006). Only one or two of rhinocerotid species is/are found: *Dicerorhinus steinheimhensis* and probably the recently-established *Plesiaceratherium balkanicum* (pers. obs.; Table 1). Data on neighboring coeval localities suggest a subtropical climate with few thermal and rain fall variations (Sabol and Kováč, 2006; Fejfar and Sabol, 2009).

**Simorre** and **Villefranche d’Astarac** refer to several localities from the same area near the eponyme villages in Gers, France (Ginsburg et al., 1975; Tassy, 1977; Antoine et al., 1997) and all dated to the early MN7/8 (~ 13 Mya). Two rhinocerotid species are distinguished: *Brachypotherium brachypus* (dominant) and *Alicornops simorrense* (Antoine et al., 1997). The environment at both localities is probably a humid forest (Bentaleb et al., 2006).

**Steinheim am Albuch** is a meteoritic crater situated 30 km North from Ulm, Baden-Württemberg, Germany (Tütken et al., 2006). It has yielded 54 mammal species including four rhinocerotids: *Alicornops simorrense, Brachypotherium brachypus, Dicerorhinus steinheimensis,* and *Lartetotherium sansaniense* (Table 1). The landscape is reconstructed as a freshwater lake surrounded by open environments under warm temperate conditions (Tütken et al., 2006; Kovar-Eder and Schweigert, 2018).

Concerning the sampling for microwear, we studied two facets – one grinding and one shearing – from the same enamel band, as illustrated in Figure 1.

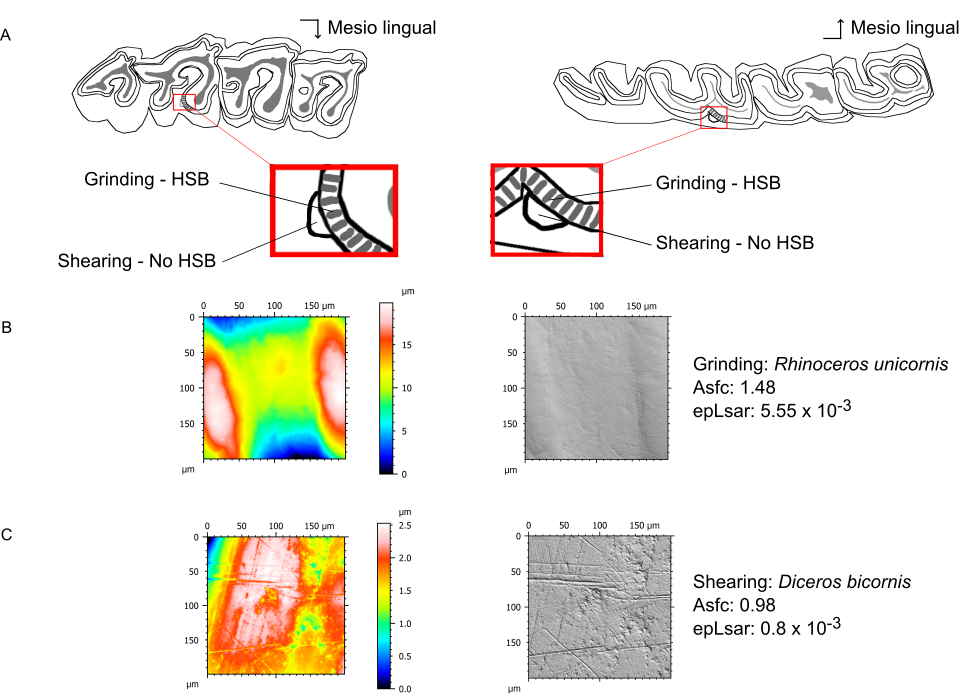

**Figure 1** **Localisation of the dental facets on rhinocerotid molars.**

Position of the two dental facets (grinding and shearing) on the second upper molar (left) and second lower molar (right). Both facets are sampled on the same enamel band with (grinding) or without (shearing) Hunter-Schreger bands (HSB). Illustration after Hullot et al. (2019).

**References**

Aiglstorfer, M., E. P. J. Heizmann, and S. Peigné. 2019. Who killed *Micromeryx flourensianus*? A case study of taphonomy and predation on ruminants in the middle Miocene of France. Lethaia 52:429–444.

Antoine, P.-O. 1997. *Aegyrcitherium beonensis* nov. gen. nov. sp., nouvel élasmothère (Mammalia, Rhinocerotidae) du gisement miocène (MN 4b) de Montréal-du-Gers (Gers, France). Position phylogénétique au sein des Elasmotheriini. Neues Jahrbuch Für Geologie Und Paläontologie - Abhandlungen 204:399–414.

Antoine, P.-O. 2002. Phylogénie et évolution des Elasmotheriina (Mammalia, Rhinocerotidae). Mémoires Du Muséum National d’Histoire Naturelle 188:5–350.

Antoine, P.-O. 2019. Rhinocerotids from the Siwalik faunal sequence; pp. in C. Badgley, D. Pilbeam, and M. Morgan (eds.), At the Foot of the Himalayas: Paleontology and Ecosystem Dynamics of the Siwalik Record of Pakistan., Johns Hopkins University Press. Baltimore.

Antoine, P.-O., and F. Duranthon. 1997. Découverte de *Protaceratherium minutum* (Mammalia, Rhinocerotidae) dans le gisement Orléanien (MN 4) de Montréal-du-Gers (Gers). Annales de Paléontologie (Vert.-Invert.) 83:201–213.

Antoine, P.-O., and J.-L. Welcomme. 2000. A New Rhinoceros From The Lower Miocene Of The Bugti Hills, Baluchistan, Pakistan: The Earliest Elasmotheriine. Palaeontology 43:795–816.

Antoine, P.-O., F. Duranthon, and P. Tassy. 1997. L’apport des grands mammifères (Rhinocérotidés, Suoidés, Proboscidiens) à la connaissance des gisements du Miocène d’Aquitaine (France). BoiChro’M97 spécial 21:581–590.

Antoine, P.-O., D. Becker, Y. Laurent, and F. Duranthon. 2018. The Early Miocene teleoceratine *Prosantorhinus* aff. *douvillei* (Mammalia, Perissodactyla, Rhinocerotidae) from Béon 2, Southwestern France. Revue de Paléobiologie 37.

Antoine, P.-O., K. F. Downing, J.-Y. Crochet, F. Duranthon, L. J. Flynn, L. Marivaux, G. Métais, A. R. Rajpar, and G. Roohi. 2010. A revision of *Aceratherium blanfordi* Lydekker, 1884 (Mammalia: Rhinocerotidae) from the Early Miocene of Pakistan: postcranials as a key. Zoological Journal of the Linnean Society 160:139–194.

Antoine, P.-O., G. Métais, M. Orliac, J. Crochet, L. Flynn, L. Marivaux, A. Rajpar, Dr. G. Roohi, and J. Welcomme. 2013. Mammalian Neogene biostratigraphy of the Sulaiman Province, Pakistan; pp. 400–422 in Fossil Mammals of Asia: Neogene Biostratigraphy and Chronology. Columbia University Press.

Becker, D., and J. Tissier. 2020. Rhinocerotidae from the early middle Miocene locality Gračanica (Bugojno Basin, Bosnia-Herzegovina). Palaeobiodiversity and Palaeoenvironments 100:395–412.

Bentaleb, I., C. Langlois, C. Martin, P. Iacumin, M. Carré, P.-O. Antoine, F. Duranthon, I. Moussa, J.-J. Jaeger, and N. Barrett. 2006. Rhinocerotid tooth enamel 18O/16O variability between 23 and 12 Ma in southwestern France. Comptes Rendus Geoscience 338:172–179.

Costeur, L., C. Guérin, and O. Maridet. 2012. Paléoécologie et paléoenvironnement du site miocène de Sansan; pp. 661–693 in S. Peigné and S. Sen (eds.), Mammifères de Sansan, Mémoires du Muséum national d’Histoire naturelle. vol. 203. Paris.

Crouzel, F., F. Duranthon, and L. Ginsburg. 1988. Découverte d’un riche gisement à petits et grands mammifères d’âge Orléanien dans le département du Gers (France). Comptes Rendus de l’Académie Des Sciences. Série 2, Mécanique, Physique, Chimie, Sciences de l’univers, Sciences de La Terre 307:101–104.

Duranthon, F., P. O. Antoine, C. Bulot, and J. P. Capdeville. 1999. Le Miocène inférieur et moyen continental du bassin d’Aquitaine Livret-guide de l’excursion des Journées Crouzel (10 et 11 juillet 1999). Bulletin de La Société d’histoire Naturelle de Toulouse 135:79–91.

Fejfar, O., and M. Sabol. 2009. Middle Miocene *Plesiodimylus* from the Devínska Nová Ves-Fissures site (western Slovakia). Bulletin of Geosciences 84:611–624.

Ginsburg, L., C. de Muizon, and P. Tassy. 1975. Découverte d’un important gisement à mastodontes dans le Miocène Moyen de Simorre (Gers). Comptes Rendus de l’Academie Des Sciences, Paris 280:1547–1549.

Göhlich, U. B., and O. Mandic. 2020. Introduction to the special issue “The drowning swamp of Gračanica (Bosnia-Herzegovina)—a diversity hotspot from the middle Miocene in the Bugojno Basin.” Palaeobiodiversity and Palaeoenvironments 100:281–293.

Heissig, K. 2012. Les Rhinocerotidae (Perissodactyla) de Sansan; pp. 317–485 in S. Peigné and S. Sen (eds.), Mammifères de Sansan. vol. 203. Mémoires du Muséum national d’Histoire naturelle, Paris.

Hullot, M., P.-O. Antoine, M. Ballatore, and G. Merceron. 2019. Dental microwear textures and dietary preferences of extant rhinoceroses (Perissodactyla, Mammalia). Mammal Research 64:397–409.

Kovar-Eder, J., and G. Schweigert. 2018. Revision of the plant assemblage of Steinheim am Albuch (Baden-Württemberg, Germany, middle Miocene, reference locality of Mammal Neogene Zone MN 7). Bulletin of Geosciences 92.

Martin, C., I. Bentaleb, and P.-O. Antoine. 2011. Pakistan mammal tooth stable isotopes show paleoclimatic and paleoenvironmental changes since the early Oligocene. Palaeogeography Palaeoclimatology Palaeoecology 311:19–29.

Métais, G., P.-O. Antoine, S. R. H. Baqri, J.-Y. Crochet, D. De Franceschi, L. Marivaux, and J.-L. Welcomme. 2009. Lithofacies, depositional environments, regional biostratigraphy and age of the Chitarwata Formation in the Bugti Hills, Balochistan, Pakistan. Journal of Asian Earth Sciences 34:154–167.

Orliac, M. J., P.-O. Antoine, and F. Duranthon. 2006. The Suoidea (Mammalia, Artiodactyla), exclusive of Listriodontinae, from the early Miocene of Béon 1 (Montréal-du-Gers, SW France, MN4). Geodiversitas 28:685–718.

Peigné, S., and S. Sen. 2012. Mammifères de Sansan, Mémoires du Muséum national d’Histoire naturelle. Muséum national d’Histoire naturelle, Paris, 709 pp.

Rage, J.-C., and S. Bailón. 2005. Amphibians and squamate reptiles from the late early Miocene (MN 4) of Béon 1 (Montréal-du-Gers, southwestern France). Geodiversitas 27:413–441.

Sabol, M., and M. Kováč. 2006. Badenian palaeoenvironment, faunal succession and biostratigraphy: a case study from northern Vienna Basin, Devínska Nová Ves-Bonanza site (Western Carpathians, Slovakia). Beiträge Zur Paläontologie 30:415–425.

Tassy, P. 1977. Découverte de *Zygolophodon turicensis* (Schinz) (Proboscidea, Mammalia) au lieu-dit Malartic à Simorre, Gers (Vindobonien moyen); implications paléoécologiques et biostratigraphiques. Geobios 10:655–669.

Tütken, T., T. W. Vennemann, H. Janz, and E. P. J. Heizmann. 2006. Palaeoenvironment and palaeoclimate of the Middle Miocene lake in the Steinheim basin, SW Germany: A reconstruction from C, O, and Sr isotopes of fossil remains. Palaeogeography, Palaeoclimatology, Palaeoecology 241:457–491.

Welcomme, J.-L., P.-O. Antoine, F. Duranthon, P. Mein, and L. Ginsburg. 1997. Nouvelles découvertes de Vertébrés miocènes dans le synclinal de Dera Bugti (Balouchistan, Pakistan). Comptes Rendus de l’Académie Des Sciences-Series IIA-Earth and Planetary Science 325:531–536.

Xafis, A., J. Saarinen, K. Bastl, D. Nagel, and F. Grímsson. 2020. Palaeodietary traits of large mammals from the middle Miocene of Gračanica (Bugojno Basin, Bosnia-Herzegovina). Palaeobiodiversity and Palaeoenvironments 100:457–477.
