## Supplementary material for "Spatio-temporal diversity of dietary preferences and stress sensibilities of early and middle Miocene Rhinocerotidae from Eurasia: Impact of climate changes": S4

setwd() #input working directories

hypo <- read.table('Hypo.txt',sep="\t",dec='.',header=T) #load your data

#Charge necessary packages

library(lme4)

library(car)

library(MASS)

#To avoid 0 in Poisson laws #

hypo$Defect <- hypo$Defect + 1

hypo$Severity <- hypo$Severity + 1

hypo$Multiple <- hypo$Multiple + 1

#For Localization, Multiple and Severity only

hypo$Defect <- factor(hypo$Defect,levels = c("2","1","3","4","5","6","8","9","10"))

\## force numerical factors to be analysed as categorical variables, change levels

hypo$Genus <- factor(hypo$Genus, levels = c("Brachypotherium","Alicornops","Dicerorhinus","Gaindatherium","Hispanotherium","Lartetotherium","Plesiaceratherium","Prosantorhinus",'Hoploaceratherium',"Mesaceratherium","Pleuroceros","Bugtirhinus","Protaceratherium"))

hypo$Tooth <- factor(hypo$Tooth,levels = c("D4","D1","D2","D3","P4","P2","P3","M1","M2","M3"))

hypo$Locality <- factor(hypo$Locality,levels = c("Beon1","Kumbi 4","Sansan","Simorre","Villefranche","Gracanica","Devinska Nova Ves","Steinheim","Beon 2"))

hypo$Species <- factor(hypo$Species,levels = c("brachypus","cf. matritense","aff. douvillei","balkanicum","beonense","blanfordi","cf. browsi","douvillei","fatehjangense","gajense","minutum",'mirallesi',"naricum","perimense","praecursor","sansaniense","simorrense","sp","steinheimensis",'tetradactylum','welcommi'))

\## get row ids for each Genus

(Genus.nms <- unique(hypo$Genus))

nGenus <- length(Genus.nms)

Genus.ids <- sapply(Genus.nms, function(x) as.numeric(rownames(subset(hypo,Genus==x))))

names(Genus.ids) <- Genus.nms

nFolds <- 5 # number of folds

oos.list <- vector('list',nFolds) # list to hold row indices for out-of-sample sets

for (i in 1:nFolds) oos.list[[i]] <- vector('list',nGenus)

for (i in 1:nGenus) {

oos.inds <- Genus.ids[[i]]

starting.n <- length(oos.inds)

mylist <- vector('list',nFolds)

for (j in 1:nFolds) {

if (j < nFolds) {

set.seed(i*1000+(j))

fold.sample <- sample(1:length(oos.inds),round(((1/nFolds)*starting.n)),replace=F)

oos.list[[j]][[i]] <- oos.inds[fold.sample]

oos.inds <- oos.inds[-fold.sample]

} else {

oos.list[[j]][[i]] <- oos.inds

}

}

}

oos <- lapply(oos.list, unlist)

sum(duplicated(unlist(oos))) # no duplicated row ids, which is what we want

training <- lapply(oos, function(x) hypo[-x,]) # list of training data frames

test <- lapply(oos, function(x) hypo[x,]) # list of test data frames

sapply(training, dim); sapply(test, dim) # dimensions OK

\## function to run full model, and perfrom cross-validation, for all models in the candidate set, for any response variable

fit.func <- function(models, response) {

\## set up data frame for storing results

res <- data.frame(model=character(),nPar=numeric(),D=numeric(),aic=numeric(),bic=numeric(),R2=numeric(),cv.R2=numeric())

fits <- vector('list',length(models))

for (i in 1:length(models)) {

model <- models[i]

model.formula <- paste(response,model,sep=' ~ ')

fit <- glmer(model.formula, data=hypo, family=binomial) # fit to full dataset

nPar <- extractAIC(fit)[1] # number of parameters

D <- -2*as.numeric(logLik(fit)) # deviance

aic <- extractAIC(fit)[2]

bic <- D + nPar*(log(nrow(hypo)))

R2 <- 1-((sum((hypo[,response]-fitted(fit))^2))/sum((hypo[,response]-mean(hypo[,response]))^2))

\## cross.validation

R2.vec <- rep(NA, nFolds) # storage vector for cross-validation R2 for each fold

for (j in 1:nFolds) {

fold.hypo <- training[[j]]

test.hypo <- test[[j]]

fold.fit <- glmer(model.formula, data=fold.hypo, family=binomial) # fit to full dataset

fold.preds <- predict(fold.fit,newdata=test.hypo,re.form=NA)

fold.R2 <- 1-((sum((test.hypo[,response]-fold.preds)^2))/sum((test.hypo[,response]-mean(test.hypo[,response]))^2))

R2.vec[j] <- fold.R2

}

cv.R2 <- mean(R2.vec)

\## store results

res <- rbind(res,data.frame(model=model,nPar=nPar,D=D,aic=aic,bic=bic,R2=R2,cv.R2=cv.R2))

fits[[i]] <- fit

print(paste('Model',i,'is done'))

}

\## sort by aic

res <- res[order(res$aic),]

\## calculate dAIC and wAIC

res$d.aic <- res$aic-res$aic[1]

exp.d.aic <- exp(-res$d.aic/2)

res$w.aic <- exp.d.aic/sum(exp.d.aic)

res <- res[,c("model","nPar","D","aic","d.aic","w.aic","bic","R2","cv.R2")]

\## name model list

names(fits) <- models

\## combine into list and return

return(list(res=res,fits=fits))

}

\## create a candidate model set - adjust this as necessary

models<-c('(1|Specimen)',

'(1|Specimen) + Genus',

'(1|Specimen) + Locality',

'(1|Specimen) + Country',

'(1|Specimen) + Age',

'(1|Specimen) + Tooth',

'(1|Specimen) + Position',

'(1|Specimen) + Side',

'(1|Specimen) + Wear')

\## run function

res <- fit.func(models=models, response='Hypo')

\## model selection table

res$res

########## refine model list above until final model selected

\## look at results for the top one

summary(res$fits$'(1|Specimen) + Tooth + Locality + Genus + Wear + Position')

##Correction for overdispersion if needed

(D <- -2*as.numeric(logLik(res$fits$'(1|Specimen) + Tooth + Locality + Genus + Wear + Position'))) # deviance

(disp <- D/df.residual(res$fits$'(1|Specimen) + Tooth + Locality + Genus + Wear + Position')) #overdispersion if disp >1

Mod1 <- glmmPQL(Hypo~ Tooth + Locality + Genus + Wear + Position, random = ~1|Specimen, family = quasibinomial,data=hypo)

summary(Mod1)

#Pairwise comparison for Species, Tooth, Locality

library(multcomp)

mc_tukey <- glht(Mod1, linfct=mcp(Tooth="Tukey"))

summary(mc_tukey) ## ## Simultaneous Tests for General Linear Hypotheses ## ## Multiple Comparisons of Means: Tukey Contrasts ## ##
