## Supplementary material for "Spatio-temporal diversity of dietary preferences and stress sensibilities of early and middle Miocene Rhinocerotidae from Eurasia: Impact of climate changes": S5

Sum up GLMM results

**Microwear**

Concerning microwear, we conducted generalized linear mixed models (GLMMs) for five response variables, corresponding to the classic DMTA parameters: Anisotropy (epLsar), Complexity (Asfc), FTfv (fine textural fill-volume), HAsfc9 and HAsfc81 (heterogeneities of the complexity).

Models were built using a modified version of the script from Arman et al. (2019). We selected Gaussian family to study our response variables. Model construction followed a bottom-up approach as described in the main text, and model selection is based on the lowest AIC score (Akaike’s Information Criterion). This resulted in 190 models were compared across the five response variables (see electronic supplementary material, S5).

Law used = Gaussian

- Variables tested: Anisotropy (epLsar), Complexity (Asfc), Heterogeneities of complexity (HAsfc9, HAsfc81), and Fine Textural fill-volume (FTfv)
- Factors: 1|Specimen (random effect), Genus, Locality, Country, Age, Tooth, Position (Upper/Lower), Side (Left/Right), Facet

References for factors:

- Genus = *Brachypotherium* because present in most localities
- Locality = Béon 1 (most abundant)
- Country = France, most localities
- Age = early Miocene, age of Béon 1
- Tooth = M2

**Anisotropy**

No over-dispersion

```
Formula: Anisotropy ~ (1 | Specimen) + Facet + Tooth

     AIC      BIC   logLik deviance df.resid
 -1301.9  -1255.4    666.9  -1333.9      119

                Estimate Std. Error t value
(Intercept)    0.0034112  0.0002917  11.695
FacetShearing  0.0005028  0.0002995   1.679
ToothM1       -0.0007043  0.0004524  -1.557
ToothM3       -0.0008982  0.0004617  -1.946
ToothM        -0.0013272  0.0006022  -2.204
```

Tooth: M3 and M have significantly lower anisotropy than M2

Pairwise comparisons did not show any other significant differences between tooth loci

Facet: no significant differences

**Complexity**

Over dispersion

```
Formula: Complexity ~ (1 | Specimen) + Locality + Genus + Side + Position +      Country + Age


     AIC      BIC   logLik deviance df.resid
   454.0    514.7   -206.0    412.0      112

                          Estimate Std. Error t value
(Intercept)                0.82464    0.35149   2.346
LocalityKumbi 4            3.04483    0.86458   3.522
LocalitySansan             1.27984    0.53037   2.413
LocalitySimorre            1.64286    0.55372   2.967
LocalityVillefranche       2.15416    0.51840   4.155
LocalityGracanica         -0.19883    0.36991  -0.538
LocalityDevinska Nova Ves  1.03554    0.83971   1.233
GenusAlicornops           -1.56915    0.51356  -3.055
GenusDicerorhinus         -0.91500    1.11073  -0.824
GenusGaindatherium        -1.21607    1.14497  -1.062
GenusHispanotherium        0.36525    0.48464   0.754
GenusLartetotherium       -0.69857    0.57585  -1.213
GenusPlesiaceratherium    -0.13553    0.36301  -0.373
GenusProsantorhinus        0.01527    0.41541   0.037
GenusHoploaceratherium     1.12791    0.60130   1.876
GenusMesaceratherium      -2.35336    0.96153  -2.448
GenusPleuroceros          -3.13030    1.13867  -2.749
SideRight                  0.50423    0.23998   2.101
PositionUpper              0.42123    0.22031   1.912
```

**Correction for over-dispersion**

```
                           Estimate Std. Error   t value  Pr(>|z|) z value
(Intercept)                0.824641   0.436143  2.346123  0.058657    1.89
LocalityKumbi 4            3.044832   1.072805  3.521740  0.004537    2.84
LocalitySansan             1.279843   0.658107  2.413097  0.051807    1.94
LocalitySimorre            1.642857   0.687073  2.966961  0.016798    2.39
LocalityVillefranche       2.154160   0.643246  4.155428  0.000811    3.35
LocalityGracanica         -0.198831   0.458999 -0.537511  0.664881   -0.43
LocalityDevinska Nova Ves  1.035539   1.041947  1.233206  0.320296    0.99
GenusAlicornops           -1.569147   0.637246 -3.055421  0.013802   -2.46
GenusDicerorhinus         -0.915000   1.378238 -0.823780  0.506760   -0.66
GenusGaindatherium        -1.216071   1.420726 -1.062094  0.392025   -0.86
GenusHispanotherium        0.365252   0.601361  0.753653  0.543602    0.61
GenusLartetotherium       -0.698570   0.714538 -1.213107  0.328246   -0.98
GenusPlesiaceratherium    -0.135531   0.450434 -0.373354  0.763499   -0.30
GenusProsantorhinus        0.015269   0.515462  0.036756  0.976369    0.03
GenusHoploaceratherium     1.127906   0.746115  1.875778  0.130609    1.51
GenusMesaceratherium      -2.353365   1.193101 -2.447521  0.048555   -1.97
GenusPleuroceros          -3.130302   1.412903 -2.749087  0.026725   -2.22
SideRight                  0.504225   0.297777  2.101104  0.090399    1.69
PositionUpper              0.421235   0.273373  1.911980  0.123346    1.54
```

Locality: Béon 1 has lower complexity than Kumbi 4, Sansan, Simorre, and Villefranche

Pairwise comparisons suggested the following differences:

- Gracanica has lower complexity than Kumbi (p-value = 0.004), Simorre (p-value = 0.027), Villefranche (p-value < 0.001)

Genus: *Brachypotherium* has a higher complexity than *Alicornops*, *Mesaceratherium*, and *Pleuroceros*

Pairwise comparisons suggested the following differences:

- *Hoploaceratherium* has higher complexity than *Alicornops* (p-value < 0.01) and *Pleuroceros* (p-value = 0.028)

Side: No significant differences when over-dispersion is corrected

Position: No significant differences when over-dispersion is corrected

**FTfv**

Important over dispersion (23.99)

```
Formula: FTFV ~ (1 | Specimen) + Facet + Country + Age
                          Estimate Std. Error t value
(Intercept)                  46550       2715  17.148
FacetShearing               -18705       3398  -5.505
CountryPakistan              20615       6159   3.347
CountryBosnia-Herzegovina   -12283       6105  -2.012
CountrySlovakia               4223       9155   0.461
AgeMiddle Miocene             5958       4016   1.484
```

**Correction for over-dispersion**

```
                           Estimate Std. Error   t value  Pr(>|z|) z value
(Intercept)                4.66e+04   5.02e+07  1.71e+01  9.99e-01       0
FacetShearing             -1.87e+04   6.29e+07 -5.50e+00  1.00e+00       0
CountryPakistan            2.06e+04   1.14e+08  3.35e+00  1.00e+00       0
CountryBosnia-Herzegovina -1.23e+04   1.13e+08 -2.01e+00  1.00e+00       0
CountrySlovakia            4.22e+03   1.69e+08  4.61e-01  1.00e+00       0
AgeMiddle Miocene          5.96e+03   7.43e+07  1.48e+00  1.00e+00       0
```

No significant differences after correction for over-dispersion

**HAfsc9**

No over-dispersion

```
Formula: H9 ~ (1 | Specimen) + Locality + Age + Country
                          Estimate Std. Error t value
(Intercept)                0.27135    0.01992  13.625
LocalityKumbi 4            0.12749    0.05045   2.527
LocalitySansan             0.18256    0.04106   4.446
LocalitySimorre            0.05836    0.06851   0.852
LocalityVillefranche       0.04936    0.05235   0.943
LocalityGracanica         -0.05625    0.04731  -1.189
LocalityDevinska Nova Ves  0.01748    0.07452   0.235
CountryPakistan            0.12749    0.05158   2.472
CountryBosnia-Herzegovina -0.17907    0.05151  -3.476
CountrySlovakia           -0.10534    0.07823  -1.347
AgeMiddle Miocene          0.12282    0.03381   3.633
```

Locality: Béon 1 has lower HAsfc9 than Kumbi 4 and Sansan

Pairwise comparisons suggested the following differences:

Sansan has higher HAsfc9 than Gracanica (p-value < 0.001)

Age: Middle Miocene rhinocerotids have higher HAsfc9 than early Miocene ones

Country: Bosnia-Herzegovina has lower HAsfc9 than France, Pakistan has higher HAsfc9 than France

**HAfsc81**

No over-dispersion

```
Formula: H81 ~ (1 | Specimen) + Locality + Age + Country

     AIC      BIC   logLik deviance df.resid
    57.6    104.0    -12.8     25.6      119

                          Estimate Std. Error t value
(Intercept)                0.53989    0.03482  15.504
LocalityKumbi 4            0.20739    0.08821   2.351
LocalitySansan             0.28099    0.07179   3.914
LocalitySimorre            0.02115    0.11979   0.177
LocalityVillefranche       0.07822    0.09153   0.855
LocalityGracanica         -0.11848    0.08272  -1.432
LocalityDevinska Nova Ves  0.06346    0.13030   0.487
CountryPakistan            0.20739    0.09191   2.256
CountryBosnia-Herzegovina -0.29705    0.09179  -3.236
CountrySlovakia           -0.11511    0.13938  -0.826
AgeMiddle Miocene          0.17857    0.06024   2.964
```

Locality: Béon 1 has a lower HAsfc81 than Kumbi 4 and Sansan

Pairwise comparisons suggested the following differences:

Gracanica has lower HAsfc81 than Sansan (p-value = 0.001)

Age: Middle Miocene rhincerotids have higher HAsfc81 than early Miocene ones

Country: French rhinocerotids have lower HAsfc81 than that from Pakistan (Kumbi 4 only) but higher than that from Bosnia-Herzegovina (Gracanica only)

Pairwise comparisons showed that Bosnia-Herzegovina (Gracanica only) had lower HAsfc81 than Pakistan (Kumbi 4 only; p-value < 0.001)

**BIC ranking differences**

Without Genus forced:

- Anisotropy, HAsfc9 and HAsfc81: best candidate model is 1|Specimen with BIC ranking
- Complexity: less factors with BIC-ranking (1|Specimen + Age)
- FTFV: less factors with BIC-ranking (1|Specimen + Facet)

**Hypoplasia**

Concerning hypoplasia, we conducted generalized linear mixed models (GLMMs) for five response variables: Hypo (presence or absence of the defect), Defect (type of defect encountered), Localization, Severity, Multiple

Models were built using a modified version of the script from Arman et al. (2019). We selected Poisson family to study our response variables except for Hypo, for which we selected Binomial law. Model construction followed a bottom-up approach as described in the main text, and model selection is based on the lowest AIC score (Akaike’s Information Criterion). This resulted in 138 models were compared across the five response variables (see electronic supplementary material, S6).

Law used = Binomial or Poisson

- Variables tested: Hypo, Defect, Localization, Multiple, Severity
- Factors: 1|Specimen (random effect), Genus, Locality, Country, Age, Tooth, Position (Upper/Lower), Side (Left/Right), Wear, Defect (converted to factor for Localization, Multiple and Severity models)

References for factors:

- Genus = *Brachypotherium* because present in most localities
- Locality = Béon 1 (most abundant)
- Country = France, most localities
- Age = early Miocene, age of Béon 1
- Tooth = D4, most commonly affected teeth in rhinos
- Wear = Average
- Defect = LEH

**Hypo**

No over-dispersion

Several final models with same AIC

```
Formula: Hypo ~ (1 | Specimen) + Tooth + Locality + Genus + Wear + Position
                           Estimate Std. Error z value Pr(>|z|)
(Intercept)                -0.07087    0.67135  -0.106 0.915932
ToothD1                    -1.46270    0.54865  -2.666 0.007676 **
ToothD2                    -3.19922    0.77300  -4.139 3.49e-05 ***
ToothD3                    -3.46584    0.77142  -4.493 7.03e-06 ***
ToothP4                    -0.55392    0.44635  -1.241 0.214604
ToothP2                    -1.64501    0.49078  -3.352 0.000803 ***
ToothP3                    -1.37676    0.47699  -2.886 0.003898 **
ToothM1                    -1.31814    0.45681  -2.886 0.003908 **
ToothM2                    -0.94897    0.44093  -2.152 0.031381 *
ToothM3                    -0.54775    0.44362  -1.235 0.216924
LocalityKumbi 4           -16.70770  119.24099  -0.140 0.888567
LocalitySansan             -3.05384    1.35564  -2.253 0.024278 *
LocalitySimorre            -0.40280    0.79999  -0.504 0.614610
LocalityVillefranche       -0.14987    0.71576  -0.209 0.834145
LocalityGracanica           1.73091    0.69947   2.475 0.013338 *
LocalityDevinska Nova Ves   0.74979    1.52741   0.491 0.623504
LocalitySteinheim           0.41462    0.95490   0.434 0.664142
LocalityBeon 2             -0.35451    1.48775  -0.238 0.811658
GenusAlicornops            -1.26368    0.51874  -2.436 0.014849 *
GenusDicerorhinus          -1.63218    1.35757  -1.202 0.229256
GenusGaindatherium         14.84370  119.24751   0.124 0.900937
GenusHispanotherium        -1.19218    0.70417  -1.693 0.090449 .
GenusLartetotherium         0.71837    1.08342   0.663 0.507291
GenusPlesiaceratherium      0.46922    0.56555   0.830 0.406722
GenusProsantorhinus         0.21457    0.58217   0.369 0.712454
GenusHoploaceratherium      2.30430    1.33709   1.723 0.084822 .
GenusMesaceratherium       14.25435  119.24371   0.120 0.904848
GenusPleuroceros           15.17913  119.24046   0.127 0.898704
GenusBugtirhinus            0.95076  266.62818   0.004 0.997155
GenusProtaceratherium       1.77357    2.79794   0.634 0.526155
WearOther                  -1.34943    1.22149  -1.105 0.269270
WearUnder                  -0.98082    0.34838  -2.815 0.004872 **
WearUpper                  -0.03947    0.23914  -0.165 0.868916
PositionUpper              -0.45626    0.23950  -1.905 0.056771 .
```

Tooth: All teeth but P4/p4 and M3/m3 were significantly less affected than D4/d4

Pairwise comparison also highlighted significant differences for the following pairs:

- D3/d3 (less affected) and D1/d1 (p-value =0.038), P4/p4 (p-value < 0.01), P2/p2 (p-value = 0.018), P3/p3 (p-value < 0.01), M1/m1 (p-value < 0.01), M2/m2 (p-value < 0.01), M3/m3 (p-value < 0.01)
- D2/d2 (less affected) and P4/p4 (p-value < 0.01), M1/m1 (p-value = 0.03), M2/m2 (p-value < 0.01), M3/m3 (p-value < 0.01)
- P4/p4 (more affected) and P2/p2 (p-value < 0.01), P3/p3 (p-value < 0.01), M1/m1 (p-value = 0.023)
- M3/m3 (more affected) and D1/d1 (p-value = 0.02), P2/p2 (p-value < 0.01), P3/p3 (p-value < 0.01), M1/m1 (p-value < 0.01)

Locality: Gračanica specimens had a higher hypoplasia prevalence than Béon 1

Pairwise comparison only found significant differences for hypoplasia prevalence between Gracanica (more affected) and Sansan (p-value = 0.011)

Genus: *Alicornops* was less prone to hypoplasia than *Brachypotherium*

Pairwise comparison also highlighted significant differences for the following pairs:

- *Plesiaceratherium* (more affected) and *Hispanotherium* (p-value = 0.02)

Wear: Less worn teeth presented less hypoplasia

Pairwise comparison also highlighted significant differences between very worn teeth (more affected) than low worn ones (p-value = 0.026)

Position: significant differences in the prevalence of hypoplasia between upper and lower teeth (upper less affected)

Tested separately (as Locality is redundant with Age and Country):

Age: no significant differences between early and middle Miocene

Country: France has significantly lower prevalence than Bosnia-Herzegovina

Tukey’s Contrasts did not reveal other significant differences between countries

**Defect**

Over-dispersion

```
Formula: Defect ~ (1 | Specimen) + Genus + Tooth + Position + Age + Wear
                        Estimate Std. Error z value Pr(>|z|)
(Intercept)             0.415144   0.173327   2.395  0.01661 *
GenusAlicornops        -0.587398   0.122219  -4.806 1.54e-06 ***
GenusDicerorhinus      -0.487421   0.175645  -2.775  0.00552 **
GenusGaindatherium     -0.314109   0.325317  -0.966  0.33427
GenusHispanotherium    -0.058239   0.169627  -0.343  0.73135
GenusLartetotherium    -0.136420   0.198787  -0.686  0.49255
GenusPlesiaceratherium  0.183916   0.136688   1.346  0.17846
GenusProsantorhinus     0.149516   0.144436   1.035  0.30059
GenusHoploaceratherium -0.188551   0.135003  -1.397  0.16252
GenusMesaceratherium   -0.424442   0.302178  -1.405  0.16014
GenusPleuroceros       -0.093292   0.210633  -0.443  0.65783
GenusBugtirhinus       -0.427167   0.563909  -0.758  0.44874
GenusProtaceratherium   1.214595   0.545589   2.226  0.02600 *
ToothD1                -0.157019   0.149676  -1.049  0.29415
ToothD2                -0.289465   0.162400  -1.782  0.07468 .
ToothD3                -0.330176   0.156044  -2.116  0.03435 *
ToothP4                 0.055138   0.127563   0.432  0.66557
ToothP2                -0.007054   0.131947  -0.053  0.95737
ToothP3                -0.011954   0.131357  -0.091  0.92749
ToothM1                -0.048694   0.127708  -0.381  0.70299
ToothM2                 0.044613   0.125056   0.357  0.72128
ToothM3                 0.267043   0.124652   2.142  0.03217 *
PositionUpper          -0.163296   0.059711  -2.735  0.00624 **
AgeMiddle Miocene       0.299140   0.138036   2.167  0.03023 *
WearOther              -0.267783   0.262869  -1.019  0.30835
WearUnder              -0.184802   0.082638  -2.236  0.02533 *
WearUpper               0.011306   0.061360   0.184  0.85381
```

**Correction for over-dispersion**

```
                            Value Std.Error  DF   t-value p-value
(Intercept)             0.4471764 0.1551884 689  2.881506  0.0041
GenusAlicornops        -0.6185213 0.1042103 672 -5.935321  0.0000
GenusDicerorhinus      -0.4464849 0.1568714 689 -2.846185  0.0046
GenusGaindatherium     -0.2909126 0.2895432 689 -1.004730  0.3154
GenusHispanotherium    -0.0559215 0.1568797 672 -0.356461  0.7216
GenusLartetotherium    -0.1335078 0.1807849 689 -0.738490  0.4605
GenusPlesiaceratherium  0.1847084 0.1270053 672  1.454336  0.1463
GenusProsantorhinus     0.1534469 0.1345922 672  1.140087  0.2547
GenusHoploaceratherium -0.2132763 0.1251805 672 -1.703749  0.0889
GenusMesaceratherium   -0.3798576 0.2722356 689 -1.395326  0.1634
GenusPleuroceros       -0.0789069 0.1918417 689 -0.411312  0.6810
GenusBugtirhinus       -0.3848068 0.4818459 689 -0.798610  0.4248
GenusProtaceratherium   1.1985500 0.5626811 689  2.130070  0.0335
ToothD1                -0.1470179 0.1258666 672 -1.168045  0.2432
ToothD2                -0.2537054 0.1347165 672 -1.883254  0.0601
ToothD3                -0.3144453 0.1277324 672 -2.461750  0.0141
ToothP4                 0.0476104 0.1081105 672  0.440386  0.6598
ToothP2                -0.0239468 0.1119175 672 -0.213968  0.8306
ToothP3                -0.0227403 0.1111812 672 -0.204534  0.8380
ToothM1                -0.0658520 0.1076980 672 -0.611451  0.5411
ToothM2                 0.0439349 0.1058307 672  0.415143  0.6782
ToothM3                 0.2950749 0.1056874 672  2.791960  0.0054
PositionUpper          -0.1604740 0.0532903 672 -3.011317  0.0027
AgeMiddle Miocene       0.3010624 0.1283901 689  2.344903  0.0193
WearOther              -0.2314999 0.2165559 672 -1.069008  0.2854
WearUnder              -0.1985132 0.0695102 672 -2.855888  0.0044
WearUpper               0.0057066 0.0521644 672  0.109397  0.9129
```

Genus: *Alicornops* and *Dicerorhinus* (less hypoplasia and/or likely to be LEH) have a different hypoplasia pattern than *Brachypotherium*

*Protaceratherium* (higher score) differs from *Brachypotherium* in the hypoplasia pattern: less likely to be simple LEH

Pairwise comparison also highlighted differences between the following pairs:

- *Alicornops* (lower score) and *Hispanotherium* (p-value = 0.064), *Plesiaceratherium* (p-value < 0.01), *Prosantorhinus* (p-value < 0.01), *Protaceratherium* (p-value = 0.05)
- *Dicerorhinus* (lower score) and *Plesiaceratherium* (p-value = 0.034), *Prosantorhinus* (p-value = 0.071)

Tooth: Different pattern of hypoplasia between D3/d3, M3/m3 and D4/d4

Pairwise comparison also highlighted differences between the following pairs:

- M3/m3 (higher score) and D1/d1 (p-value < 0.01), D2/d2 (p-value < 0.01), D3/d3 (p-value < 0.01), P4/p4 (p-value = 0.013), P2/p2 (p-value < 0.01), P3/p3 (p-value < 0.01), M1/m1 (p-value < 0.01), M2/m2 (p-value < 0.01)
- D3/d3 (lower score) and P4/p4 (p-value = 0.044), M2/m2 (p-value = 0.043)

Position: Upper teeth have less defect and/or different pattern

Age: Middle Miocene specimens have more hypoplasia and/or different defect than early Miocene ones

Wear: Less worn teeth have less defect and/or different pattern

Pairwise comparison also highlighted significant differences between very worn teeth (higher score) than low worn ones (p-value = 0.031)

**Multiple**

Over-dispersion

```
Formula: Multiple ~ (1 | Specimen) + Defect + Genus
                        Estimate Std. Error z value Pr(>|z|)
(Intercept)             0.890675   0.082373  10.813  < 2e-16 ***
Defect1                -0.892273   0.062557 -14.263  < 2e-16 ***
Defect3                -0.175345   0.110994  -1.580   0.1142
Defect4                -0.032124   0.160461  -0.200   0.8413
Defect5                 0.471244   0.107940   4.366 1.27e-05 ***
Defect6                 0.193636   0.264136   0.733   0.4635
Defect8                 0.508737   0.305162   1.667   0.0955 .
Defect9                -0.179827   0.120417  -1.493   0.1353
Defect10                0.205414   0.239554   0.857   0.3912
GenusAlicornops        -0.022684   0.108181  -0.210   0.8339
GenusDicerorhinus      -0.026599   0.140961  -0.189   0.8503
GenusGaindatherium     -0.001605   0.266985  -0.006   0.9952
GenusHispanotherium    -0.015103   0.116404  -0.130   0.8968
GenusLartetotherium    -0.032663   0.180324  -0.181   0.8563
GenusPlesiaceratherium  0.024178   0.079272   0.305   0.7604
GenusProsantorhinus     0.007664   0.079800   0.096   0.9235
GenusHoploaceratherium -0.008422   0.117774  -0.072   0.9430
GenusMesaceratherium    0.028594   0.228555   0.125   0.9004
GenusPleuroceros       -0.009157   0.161563  -0.057   0.9548
GenusBugtirhinus        0.001599   0.504768   0.003   0.9975
GenusProtaceratherium   0.002292   0.532077   0.004   0.9966
```

Correction for over-dispersion

```
Fixed effects: Multiple ~ Defect + Genus
                            Value  Std.Error  DF   t-value p-value
(Intercept)             0.8961173 0.01764498 690  50.78597  0.0000
Defect1                -0.8898299 0.01171551 677 -75.95315  0.0000
Defect3                -0.1824403 0.02087744 677  -8.73863  0.0000
Defect4                -0.0587308 0.03001289 677  -1.95685  0.0508
Defect5                 0.3746201 0.02122356 677  17.65115  0.0000
Defect6                 0.2631510 0.05664021 677   4.64601  0.0000
Defect8                 0.5585553 0.05852955 677   9.54313  0.0000
Defect9                -0.1894110 0.02232116 677  -8.48571  0.0000
Defect10                0.1840063 0.04607493 677   3.99363  0.0001
GenusAlicornops        -0.0402844 0.02187716 677  -1.84139  0.0660
GenusDicerorhinus      -0.0331278 0.03188497 690  -1.03898  0.2992
GenusGaindatherium     -0.0080727 0.05590544 690  -0.14440  0.8852
GenusHispanotherium    -0.0107051 0.02619529 677  -0.40867  0.6829
GenusLartetotherium    -0.0318401 0.03902706 690  -0.81585  0.4149
GenusPlesiaceratherium  0.0078078 0.01886163 677   0.41395  0.6790
GenusProsantorhinus     0.0043298 0.01912621 677   0.22638  0.8210
GenusHoploaceratherium -0.0162640 0.02648683 677  -0.61404  0.5394
GenusMesaceratherium    0.0182583 0.05088785 690   0.35879  0.7199
GenusPleuroceros       -0.0164917 0.03477050 690  -0.47430  0.6354
GenusBugtirhinus       -0.0062874 0.09169759 690  -0.06857  0.9454
GenusProtaceratherium   0.0122369 0.14001666 690   0.08740  0.9304
```

Defect: Absence of defect (Defect1), Pits (Defect3), Aplasia (Defect4), and other defects (Defect9) less likely to be multiple than LEH

All other defects multiple by definition (combination of defects, more likely to be multiple than

All pairwise comparison significant (p-value < 0.001) except:

- Aplasia vs LEH
- Other vs Pits
- LEH + Pits and LEH + Aplasia, All
- LEH + Aplasia and LEH + Other

Genus

*Alicornops* less likely to have multiple defects than *Brachypotherium*

Tukey’s Contrasts did not reveal other significant differences between genera

**Localization**

No over-dispersion

```
Formula: Localization ~ (1 | Specimen) + Defect + Position
                Estimate Std. Error z value Pr(>|z|)
(Intercept)    2.128e-01  8.925e-02   2.384 0.017121 *
Defect1       -4.727e+01  2.025e+06   0.000 0.999981
Defect3        2.430e-01  1.237e-01   1.964 0.049548 *
Defect4       -1.308e-01  2.056e-01  -0.636 0.524587
Defect5        1.702e-01  1.572e-01   1.083 0.278973
Defect6       -2.068e-01  3.861e-01  -0.536 0.592179
Defect8        1.499e-01  4.158e-01   0.360 0.718508
Defect9        8.942e-02  1.449e-01   0.617 0.537064
Defect10      -1.166e-01  3.404e-01  -0.343 0.731843
PositionUpper  3.305e-01  9.639e-02   3.429 0.000606 ***
```

Position: Upper teeth display defects less likely to be positioned labially

Defect: Defect3 (Pits) are less likely to be on labial side

Pairwise comparisons highlighted differences between the following pairs:

- LEH and no defect (p-value < 0.001), pits (p-value < 0.001), aplasia (p-value < 0.001), LEH + Pits (p-value < 0.001), LEH + Aplasia (p-value < 0.001)

```
              Estimate Std. Error z value Pr(>|z|)
1 - 2 == 0  -3.359e+01  3.516e+04  -0.001  1.00000
3 - 2 == 0   3.345e-01  3.151e-02  10.613  < 0.001 ***
4 - 2 == 0  -2.721e-01  4.517e-02  -6.023  < 0.001 ***
5 - 2 == 0   2.049e-01  3.274e-02   6.259  < 0.001 ***
6 - 2 == 0   4.882e-01  8.616e-02   5.666  < 0.001 ***
8 - 2 == 0   1.637e-02  7.848e-02   0.209  1.00000
9 - 2 == 0   1.568e-01  3.874e-02   4.048  0.00119 **
10 - 2 == 0  2.933e-01  6.982e-02   4.201  < 0.001 ***
3 - 1 == 0   3.393e+01  3.516e+04   0.001  1.00000
4 - 1 == 0   3.332e+01  3.516e+04   0.001  1.00000
5 - 1 == 0   3.380e+01  3.516e+04   0.001  1.00000
6 - 1 == 0   3.408e+01  3.516e+04   0.001  1.00000
8 - 1 == 0   3.361e+01  3.516e+04   0.001  1.00000
9 - 1 == 0   3.375e+01  3.516e+04   0.001  1.00000
10 - 1 == 0  3.389e+01  3.516e+04   0.001  1.00000
4 - 3 == 0  -6.065e-01  4.949e-02 -12.256  < 0.001 ***
5 - 3 == 0  -1.295e-01  3.646e-02  -3.552  0.00764 **
6 - 3 == 0   1.537e-01  9.105e-02   1.688  0.68387
8 - 3 == 0  -3.181e-01  8.434e-02  -3.771  0.00354 **
9 - 3 == 0  -1.777e-01  4.630e-02  -3.837  0.00287 **
10 - 3 == 0 -4.111e-02  6.676e-02  -0.616  0.99918
5 - 4 == 0   4.770e-01  5.051e-02   9.443  < 0.001 ***
6 - 4 == 0   7.602e-01  9.669e-02   7.862  < 0.001 ***
8 - 4 == 0   2.884e-01  9.035e-02   3.192  0.02627 *
9 - 4 == 0   4.289e-01  5.312e-02   8.073  < 0.001 ***
10 - 4 == 0  5.654e-01  7.468e-02   7.571  < 0.001 ***
6 - 5 == 0   2.832e-01  9.157e-02   3.093  0.03581 *
8 - 5 == 0  -1.886e-01  8.484e-02  -2.223  0.31499
9 - 5 == 0  -4.814e-02  4.845e-02  -0.994  0.97865
10 - 5 == 0  8.841e-02  7.199e-02   1.228  0.92673
8 - 6 == 0  -4.718e-01  1.164e-01  -4.053  0.00125 **
9 - 6 == 0  -3.314e-01  9.447e-02  -3.508  0.00911 **
10 - 6 == 0 -1.948e-01  1.102e-01  -1.768  0.62839
9 - 8 == 0   1.404e-01  8.752e-02   1.605  0.73991
10 - 8 == 0  2.770e-01  1.048e-01   2.642  0.12564
10 - 9 == 0  1.366e-01  7.703e-02   1.773  0.62419
```

**Severity**

Over-dispersion

```
Formula: Severity ~ (1 | Specimen) + Defect + Genus
                        Estimate Std. Error z value Pr(>|z|)
(Intercept)             1.262300   0.072284  17.463  < 2e-16 ***
Defect1                -1.241933   0.055449 -22.398  < 2e-16 ***
Defect3                -0.097483   0.090750  -1.074  0.28274
Defect4                 0.320150   0.117166   2.732  0.00629 **
Defect5                 0.080080   0.106281   0.753  0.45117
Defect6                 0.287533   0.213837   1.345  0.17874
Defect8                 0.061406   0.314877   0.195  0.84538
Defect9                 0.024796   0.093854   0.264  0.79163
Defect10                0.071042   0.214558   0.331  0.74056
GenusAlicornops        -0.030760   0.100828  -0.305  0.76031
GenusDicerorhinus      -0.008549   0.131182  -0.065  0.94804
GenusGaindatherium     -0.029123   0.257596  -0.113  0.90999
GenusHispanotherium    -0.048333   0.109491  -0.441  0.65890
GenusLartetotherium    -0.035578   0.162097  -0.219  0.82627
GenusPlesiaceratherium -0.008138   0.071862  -0.113  0.90984
GenusProsantorhinus    -0.034339   0.072650  -0.473  0.63645
GenusHoploaceratherium -0.044668   0.109514  -0.408  0.68337
GenusMesaceratherium    0.048830   0.217563   0.224  0.82242
GenusPleuroceros        0.021749   0.150157   0.145  0.88484
GenusBugtirhinus       -0.020367   0.504072  -0.040  0.96777
GenusProtaceratherium   0.220174   0.442579   0.497  0.61885


Correction for over-dispersion
                           Value  Std.Error  DF    t-value p-value
(Intercept)             1.2645725 0.01653998 690   76.45552  0.0000
Defect1                -1.2487685 0.01055530 677 -118.30726  0.0000
Defect3                -0.1202594 0.01818718 677   -6.61231  0.0000
Defect4                 0.2925249 0.02416442 677   12.10561  0.0000
Defect5                 0.0357413 0.02001734 677    1.78552  0.0746
Defect6                 0.3081339 0.04820506 677    6.39215  0.0000
Defect8                 0.0043175 0.05903540 677    0.07313  0.9417
Defect9                 0.0045700 0.01863420 677    0.24525  0.8063
Defect10                0.0589471 0.04040143 677    1.45903  0.1450
GenusAlicornops        -0.0769105 0.02017245 677   -3.81265  0.0002
GenusDicerorhinus      -0.0066774 0.03069128 690   -0.21757  0.8278
GenusGaindatherium     -0.0200343 0.05444654 690   -0.36796  0.7130
GenusHispanotherium    -0.0467153 0.02519646 677   -1.85404  0.0642
GenusLartetotherium    -0.0278181 0.03734524 690   -0.74489  0.4566
GenusPlesiaceratherium -0.0125438 0.01812874 677   -0.69193  0.4892
GenusProsantorhinus    -0.0268300 0.01843101 677   -1.45570  0.1459
GenusHoploaceratherium -0.0234050 0.02562584 677   -0.91334  0.3614
GenusMesaceratherium    0.0315409 0.04956265 690    0.63639  0.5247
GenusPleuroceros        0.0159752 0.03357846 690    0.47576  0.6344
GenusBugtirhinus       -0.0158039 0.09001510 690   -0.17557  0.8607
GenusProtaceratherium   0.2288789 0.13251154 690    1.72724  0.0846

Defect: Absence of defect (Defect1) and Pits (Defect3) less severe than LEH
Aplasia (Defect4) is more severe than LEH
LEH + Aplasia (Defect6), LEH + Pit (Defect 5) more severe than LEH

Pairwise comparison

               Estimate Std. Error  z value Pr(>|z|)
1 - 2 == 0  -1.2487685  0.0104751 -119.213  < 0.001 ***
3 - 2 == 0  -0.1202594  0.0180491   -6.663  < 0.001 ***
4 - 2 == 0   0.2925249  0.0239809   12.198  < 0.001 ***
5 - 2 == 0   0.0357413  0.0198653    1.799  0.62459
6 - 2 == 0   0.3081339  0.0478390    6.441  < 0.001 ***
8 - 2 == 0   0.0043175  0.0585871    0.074  1.00000
9 - 2 == 0   0.0045700  0.0184927    0.247  1.00000
10 - 2 == 0  0.0589471  0.0400946    1.470  0.83413
3 - 1 == 0   1.1285092  0.0167224   67.485  < 0.001 ***
4 - 1 == 0   1.5412934  0.0231960   66.446  < 0.001 ***
5 - 1 == 0   1.2845098  0.0193419   66.411  < 0.001 ***
6 - 1 == 0   1.5569024  0.0480746   32.385  < 0.001 ***
8 - 1 == 0   1.2530861  0.0584176   21.450  < 0.001 ***
9 - 1 == 0   1.2533386  0.0168088   74.565  < 0.001 ***
10 - 1 == 0  1.3077156  0.0393674   33.218  < 0.001 ***
4 - 3 == 0   0.4127842  0.0272852   15.128  < 0.001 ***
5 - 3 == 0   0.1560006  0.0229670    6.792  < 0.001 ***
6 - 3 == 0   0.4283932  0.0503219    8.513  < 0.001 ***
8 - 3 == 0   0.1245769  0.0604467    2.061  0.43775
9 - 3 == 0   0.1248294  0.0227390    5.490  < 0.001 ***
10 - 3 == 0  0.1792064  0.0400919    4.470  < 0.001 ***
5 - 4 == 0  -0.2567836  0.0281837   -9.111  < 0.001 ***
6 - 4 == 0   0.0156090  0.0526300    0.297  1.00000
8 - 4 == 0  -0.2882073  0.0625163   -4.610  < 0.001 ***
9 - 4 == 0  -0.2879549  0.0275769  -10.442  < 0.001 ***
10 - 4 == 0 -0.2335778  0.0437707   -5.336  < 0.001 ***
6 - 5 == 0   0.2723926  0.0510122    5.340  < 0.001 ***
8 - 5 == 0  -0.0314237  0.0612097   -0.513  0.99982
9 - 5 == 0  -0.0311713  0.0246500   -1.265  0.92214
10 - 5 == 0  0.0232058  0.0421899    0.550  0.99969
8 - 6 == 0  -0.3038163  0.0752905   -4.035  0.00150 **
9 - 6 == 0  -0.3035639  0.0505042   -6.011  < 0.001 ***
10 - 6 == 0 -0.2491868  0.0617429   -4.036  0.00136 **
9 - 8 == 0   0.0002525  0.0605891    0.004  1.00000
10 - 8 == 0  0.0546295  0.0702478    0.778  0.99627
10 - 9 == 0  0.0543770  0.0422272    1.288  0.91417
```

Genus

*Alicornops* have less severe hypoplasia than *Brachypotherium*

Tukey’s Contrasts did not reveal other significant differences between genera

**BIC ranking differences**

Without Genus forced:

- Hypo: less factors in BIC-ranking selection (1|Specimen + Country)
- Defect: best candidate model 1|Specimen with BIC
- Severity and Multiple: less factors in BIC-ranking selection (1|Specimen + Defect)
- Localization same model
